# Cilium induction triggers differentiation of glioma stem cells

**DOI:** 10.1101/2020.12.23.424140

**Authors:** Gladiola Goranci-Buzhala, Aruljothi Mariappan, Lucia Ricci-Vitiani, Natasa Josipovic, Simone Pacion, Marco Gottardo, Johannes Ptok, Giuliano Callaini, Krishnaraj Rajalingam, Brian Dynlacht, Kamyar Hadian, Argyris Papantonis, Roberto Pallini, Jay Gopalakrishnan

## Abstract

Glioblastoma multiforme (GBM) possesses glioma stem cells (GSCs) that promote self-renewal, tumor propagation, and relapse. GBM has a poor prognosis, and currently, there are no curative options exist. Understanding the mechanisms of GSCs self-renewal can offer targeted therapeutic interventions. However, insufficient knowledge of the fundamental biology of GSCs is a significant bottleneck hindering these efforts. Here, we show that patient-derived GSCs recruit an elevated level of proteins that ensure the temporal cilium disassembly, leading to suppressed ciliogenesis. Depleting the cilia disassembly complex components at the ciliary base is sufficient to induce ciliogenesis in a subset of GSCs. Importantly, restoring ciliogenesis caused GSCs to behave like healthy NPCs switching from self-renewal to differentiation. Finally, using an organoid-based glioma invasion assay and brain xenografts in mice, we establish that ciliogenesis-induced differentiation can prevent the infiltration of GSCs into the brain. Our findings illustrate a crucial role for cilium as a molecular switch in determining GSCs’ fate and suggest that cilium induction is an attractive strategy to intervene in GSCs proliferation.

## Introduction

Glioblastoma (GBM) is the most frequent malignant primary brain tumor (Matsukado et al., 1961; Ferreri et al., 2010; Ostrom et al., 2014). Despite surgical resection, its rapid growth, resistance to chemotherapy, and high invasiveness cause GBM patients succumb to the disease with a median survival time of 15 months (Johnson and O’Neill, 2012; Stupp et al., 2005). Low passage patient-derived glioma stem cells (GSCs) are phenotypically similar to *in vivo* tumors characterized by their self-renewal and multi-lineage differentiation (Jacob et al., 2020; Pine et al., 2020; Singh et al., 2003; Wang et al., 2019; Lathia et al., 2015). These cell types are responsible for most aspects of tumor initiation, maintenance, and invasion *in vivo*, and thus, GSCs are the accepted models of GBM biology (Liau et al., 2017; Suvà et al., 2014). GSCs possess neural stem cell attributes exhibiting uncontrolled self-renewal properties. This could be due to genetic alterations in GSCs, adaptation between proliferative and slow-cycling states, and enrichment of stemness upon therapy (Liau et al., 2017; Neftel et al., 2019; Park et al., 2017; Rajakulendran et al., 2019; Ricci-Vitiani et al., 2010; Wang et al., 2019). However, it remains unclear whether there are also alterations in cellular structures implicated in cell cycle control, self-renewal, and differentiation of GSCs. Intense efforts are being made to understand the mechanisms of GSCs proliferation that can be exploited for therapeutic interventions.

The primary cilium is a microtubule-based cellular structure in which the minus end of the ciliary microtubule is anchored to a basal body that serves as a template for the assembly of the ciliary microtubule **(Figure S1A)** (Anvarian et al., 2019; Larsen et al., 2013). The cilium serves as a ‘cellular antenna’ sensing multiple signals, including sonic hedgehog (Shh), G-protein coupled receptors, and receptor tyrosine kinase (Nachury and Mick, 2019; Schmidt et al., 2002; Schou et al., 2015). In cycling cells, the cilium assembly occurs during cell cycle exit (G1-G0), and disassembly coincides with cell cycle re-entry (G1-S to M) (Jackson, 2011). A delay or failure in cilium disassembly could act as a brake, retaining cells in G0/G1 and transiently preventing cell cycle progression. This provides a conceptually novel ‘cilium checkpoint’ regulating cell cycle progression **(Figure S1B)**. Studies have recently begun to uncover the mechanisms linking cilia disassembly to cell cycle re-entry and identified mitotic kinases such as Aurora-A, Plk1, and Nek2 that can trigger cilium disassembly (Kim et al., 2011; Pugacheva et al., 2007; Wang et al., 2013). However, whether these protein components are assembled as a cytoplasmic complex, presumably as a cilia disassembly complex (CDC), has remained unknown for a precise temporal cilia disassembly.

CPAP, a conserved centrosomal protein, provides a scaffold for recruiting CDC components of Nde1, HDAC6, Aurora-A, and OFD1, to the ciliary base that is critical for temporal cilia disassembly at the onset of cell cycle re-entry (Gabriel et al., 2016). The precise timing of cilia disassembly ensures the length of G1-S transition and, thus, the self-renewal property of NPCs (Gabriel et al., 2016; Li et al., 2011). Prolonging the G1 phase due to a delayed cilia disassembly is sufficient to cause premature differentiation of NPCs into early neurons, leading to an overall reduction in the NPCs pool (Gabriel et al., 2016). In summary, these studies have successfully modeled the critical function of cilia dynamics in NPC maintenance.

Interestingly, frequent loss of cilia is common in various types of cancers, including breast, prostate, skin, melanoma, and pancreatic tumors (Fabbri et al., 2019; Seeley and Nachury, 2009; Zingg et al., 2018). So far, only a few reports have characterized loss of cilium in cultured GBM cells raising the question of whether cilia loss is due to cell culture artifacts, suppressed ciliogenesis, or perturbation of normal cilium formation (Moser et al., 2009; Yang et al., 2013). Although the loss of cilia in GBM has been reported, only Moser and colleagues have conducted an ultrastructural study using GBM tumors (Moser et al., 2014). Their study characterized that disruptions occur at the early stages of ciliogenesis, questioning the identity of candidate proteins that suppress the early stages of ciliogenesis (Moser et al., 2009). From this, we speculated that patient-derived GSCs have suppressed ciliogenesis at the early stage of ciliogenesis and, CDC proteins are involved in this critical period to suppress ciliogenesis. This may provide a selective advantage to GSCs to continuously self-renew without cell cycle exit and differentiation **(Figure S1B right panel).** If so, we hypothesize that perturbing these CDC components could promote ciliogenesis, and as a result, this may impair the self-renewal of GSCs and trigger them to differentiate **(Figure S1C).**

In this work, we show that irrespective of various cellular state of GBM, a panel of tested patient derived GSCs and clinical glioma tissues exhibit suppressed ciliogenesis due to elevated levels of CDC recruitment to basal bodies. We then establish that stopping CDC recruitment and altering their dynamic localization behavior at the basal body is sufficient to induce cilium in a subset of GSCs that overexpress constitutively active receptor tyrosine kinase PDGFR-α. Upon cilium induction, PDGFR-α is sequestered to newly induced cilium from its original location with concomitant reduction of overall PDGFR-α levels. Inducing ciliogenesis triggers GSCs switching from self-renewal to differentiation state. Finally, we demonstrate that GSCs induced with cilia failed to infiltrate into iPSC-derived human brain organoids and mouse brain, suggesting that cilium induction can play an instructive role in determining GSCs’ fate.

## Results

### Patient-derived GSCs exhibit suppressed ciliogenesis

To test if low-passage GSCs display suppressed ciliogenesis, we investigated at least seven patient-derived GSC lines and nine clinical glioblastoma tissues. First, we aimed to categorize the cellular states of GSC lines. A recent report indicated that GBM cells exist within four cellular states of neural-progenitor-like (NPC-like), oligodendrocyte-progenitor-like (OPC-like), astrocyte-like (AC-like), and mesenchymal-like (MES-like). Interestingly, each cellular state associates with a distinct signature of genetic alterations occurring in CDK4 (NPCs-like), PDGFRA (OPC-like), EGFR (AC-like), and NF1 (MES-like) (Neftel et al., 2019). To analyze the coexistence of these cellular states in our primary GSC cultures, we quantified relative gene expressions of CDK4, PDGFRA, EGFR, and NF1 and categorized them as NPCs-like, OPC-like, AC-like, and MES-like states. We noticed that although each of these states coexists in an individual GSC line, one state is predominantly existing over the other three in any GSC line we tested **(Figure 1A).** Thus, by measuring these gene’s relative expression levels, we categorized our patient-derived GSCs according to the known cellular states of GBM **(Figure 1A)**. We then determined that GSCs display neural stem cell markers similar to induced Pluripotent Stem Cells (iPSCs)-derived neural progenitor cells (NPCs) **(Figure S2A)**. Scoring cilia, we noticed that only a fraction of GSCs displayed cilia suggesting that suppressed ciliogenesis a common denominator irrespective of the cellular status. Like cultured GSCs, WHO grade IV GBM tissues also revealed reduced frequencies of ciliated cells **(Figure 1B-C and S2B-C).**

**Figure 1.**
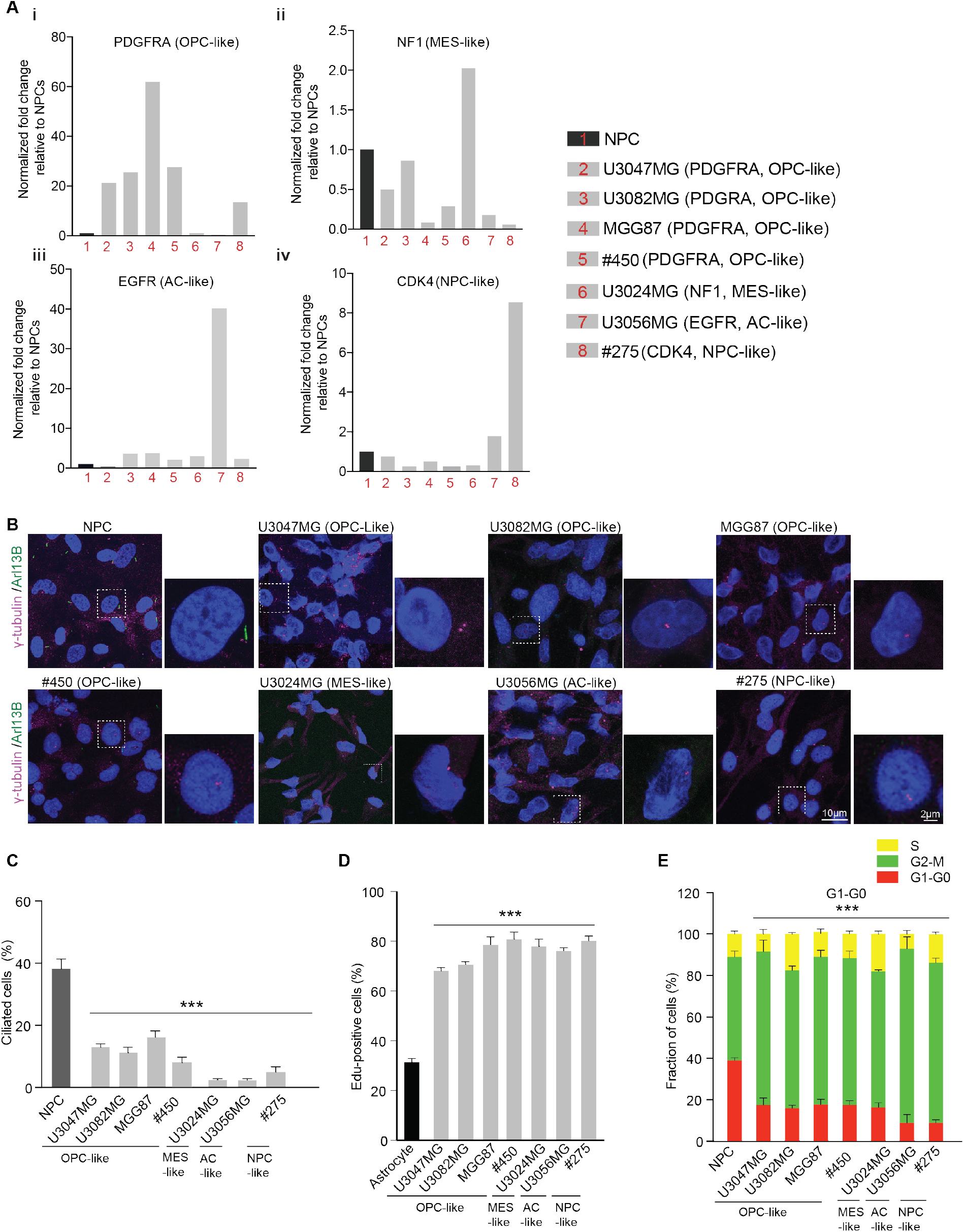
Patient-derived GSCs exhibit suppressed ciliogenesis. **A.** qRT-PCR analysis of *PDGFRA* (i), *NF1* (ii), *EGFR* (iii) and *CDK4* (iv) relative to NPCs reveal various cellular states. The calculated values were normalized to *GAPDH* expression and the values given are relative to healthy NPCs. The values are mean from three technical replicates from n=2 independent experiments. **B.** Compared to iPSCs-derived NPCs, GSCs of different cellular statuses exhibit suppressed ciliogenesis. Arl13B (green) labels cilia and γ-tubulin (magenta) labels centrosomes. Scale bar, 10 μm (overview), 2 μm (inset). At least 500 cells for each cell line from n=4 independent experiments. **C.** The bar diagram quantifies frequencies of ciliated cells. Note that compared to NPCs, GSCs of different cellular statuses displays an overall decrease in ciliation. At least 500 cells in each cell line from three (n=3) independent experiments. One-way ANOVA followed by Dunnett’s multiple comparisons test ***P <0.0001. Error bars show +/− SEM. **D.** The bar diagram quantifies frequencies of EdU positive cells. Note that GSCs exhibit an increased EdU incorporation compared to ciliated cycling astrocytes. At least, 300 cells in each cell line from three (n=3) independent experiments. One-way ANOVA, followed by Dunnett’s multiple comparisons test, ***P<0.0001. Error bars show +/− SEM. **E.** The bar diagram quantifies cell cycle analysis by Fluorescence Ubiquitin Cell Cycle Indicator (FUCCI) tracing. Note that a significant fraction of GSCs and NPCs are retained at G2-M (green). At least 100 cells in each cell line from (n=4) independent experiments. Ordinary two-way ANOVA, followed by Tukey’s multiple comparisons test, ***P <0.0001. Error bars show +/− SEM

Delayed cilia disassembly is associated with suppression of cell division via an extended G1-S transition while an accelerated cilia disassembly or loss of cilia promoted cell proliferation (Gabriel et al., 2016; Kim et al., 2011; Li et al., 2011; Pugacheva et al., 2007). As ciliogenesis is suppressed in all tested GSCs, we speculated that these GSCs would exhibit an increased proliferation. Our 24-hrs pulse labeling using ethynyl-deoxyuridine (EdU) revealed that in contrast to proliferating ciliated astrocytes, an increased number of GSCs with EdU incorporation was observed **(Figure 1D).** Furthermore, our Fluorescence Ubiquitination Cell Cycle Indicator (FUCCI)-based analyses revealed that similar to fast proliferating NPCs, only a small fraction of GSCs reside at G1/G0, a cell cycle stage at which the cilium assembly occurs in healthy cells **(Figure 1E).** These data suggest that suppressed ciliogenesis in GSCs is associated with increased proliferation.

### GSCs have an elevated level of CDC components

Because cilia formation is suppressed in GSCs at the early stages of ciliogenesis, we speculated that CDC protein levels are elevated in this critical period to suppress ciliogenesis, allowing cells to proliferate continuously. Studies have shown that Nek2, an S/G2 kinase, is elevated in cancer cells and plays a prominent role in cilia disassembly by activating its physiological substrate Kif24, a depolymerizing microtubule kinesin that suppresses cilia formation (Kim et al., 2015). By analyzing purified FLAG-tagged CPAP complexes from HEK cell extracts, we first identified that Nek2 co-purifies with the known components of the CDC. Reciprocally, CDC components of Aurora-A, HDAC6, Nde-1, OFD1, and CPAP co-purify with FLAG-tagged Nek2 complexes, which included Kif24 **(Figure S3A)**. To test our speculation that GSCs exhibit an overall increase in CDC protein levels, we performed a semi-quantitative Western blot analysis by probing an equivalent quantity of protein extracts. GSCs extracts showed an overall increase in CDC components compared to ciliated NPCs **(Figure 2A and B).** Interestingly, we measured the highest rise of Nek2 in OPC-like GSCs. Finally, immunostaining of GSC lines using specific antibodies further revealed that compared to NPCs, basal bodies of GSCs that exhibit suppressed ciliogenesis recruit an enhanced level of these proteins **(Figure 2C-D).** Together, these data suggest that CDC components are possible candidate proteins whose elevated levels could suppress ciliogenesis in tested GSCs.

**Figure 2.**
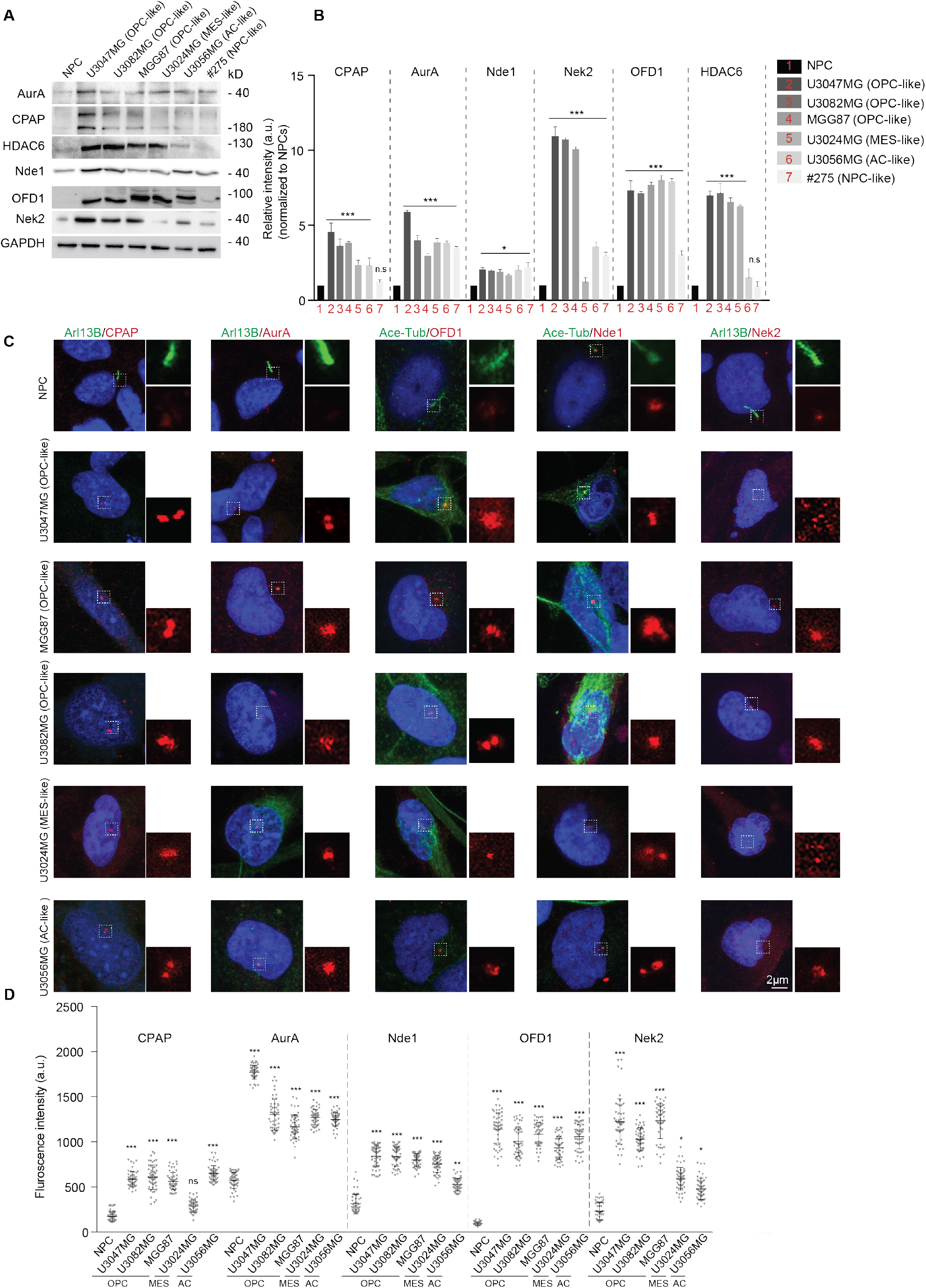
GSCs have elevated levels of CDC components. **A.** Semi-quantitative Western blot exhibiting relative levels of tested CDC components between healthy NPCs and GSCs of different cellular statuses. An equivalent quantity of cell extracts was loaded. GSCs show an overall elevated level of CDC components. GAPDH was used as a loading control. **B.** The bar diagram quantifies the relative intensity of tested CDC components in healthy NPCs and GSCs. Western blots from three (n=3) independent experiments. Ordinary two-way ANOVA followed by Sidak’s multiple comparisons test, ***P<0.0001, * P<0.01, ns represents non-significant. Error bars show +/− SEM. **C.** Compared to NPCs, GSCs of different cellular statuses recruit an enhanced level of CDC components at the ciliary base. Cilia are labeled either by Arl13B or acetylated α-tubulin (green). CDC components (red) were immunostained using antibodies specific to CPAP, Aurora-A, OFD1, Nde1 and Nek2. Scale bar 2 μm. **D.** Bar diagram quantifies fluorescence signal intensity. At least 200 cells in each cell line from (n=4) independent experiments. Ordinary one–way ANOVA followed by Tukey’s multiple comparisons test. ***P <0.0001, **P <0.001, *P <0.01, ns represents non-significant. Error bars show +/− SEM.

### Suppressing CDC level induces cilia in GSCs

First, we developed a proof-of-principle assay by testing if shRNA-mediated depletion of CDC components can induce cilia in U3047MG cultures. We detected the highest increase of ciliated cells in Nek2 depleted cultures **(Figure S3B).** We chose to use Nek2 as a tool in our experiments for a few reasons. First, due to its relatively smaller size, we were able to handle the construct in multiple experiments. Second, Nek2 but not catalytically inactive Nek2 (Nek2-KD, Nek2-kinase-dead where lysine is replaced with arginine, Nek2-K37R) can phosphorylate its physiological substrate Kif24, a kinesin that depolymerizes ciliary microtubule (Kim et al., 2015; Fry et al., 1995; Kobayashi et al., 2011). Therefore, instead of depleting the endogenous Nek2 in our experiments, we could express both catalytically active and inactive versions in a controlled manner. To test if catalytically inactive Nek2 (Nek2-KD) is sufficient to induce cilia in GSCs, we generated GSC lines that were engineered to express green fluorescence protein (GFP)-tagged Nek2-WT and catalytically inactive Nek2-KD upon doxycycline induction. The presence of GFP recognized the expressions of these inducible Nek2 variants.

In contrast to naïve or Nek2-WT expressing U3047MG GSCs, a significant number of Nek2-KD expressing GSCs displayed cilia within 24-hrs after doxycycline-induction **(Figure 3A-B)**. The frequencies of ciliated GSCs did not vary significantly after 72-hrs of doxycycline-induction (data not shown). Interestingly, Nek2-KD expression could induce cilia only in OPC-like GSC lines which exhibit the highest level of Nek2 expression (responder lines such as U3047MG, U3082MG, and MGG87) but not in other subsets of GSC lines (non-responder lines such as MES-, AC- and NPC-like GSCs) **(Figure 3C and S3C).** Cell proliferation analysis determined by Edu incorporation and Ki67 staining indicated that responder lines proliferated significantly slower than non-responder lines **(Figure 3D).**

**Figure 3.**
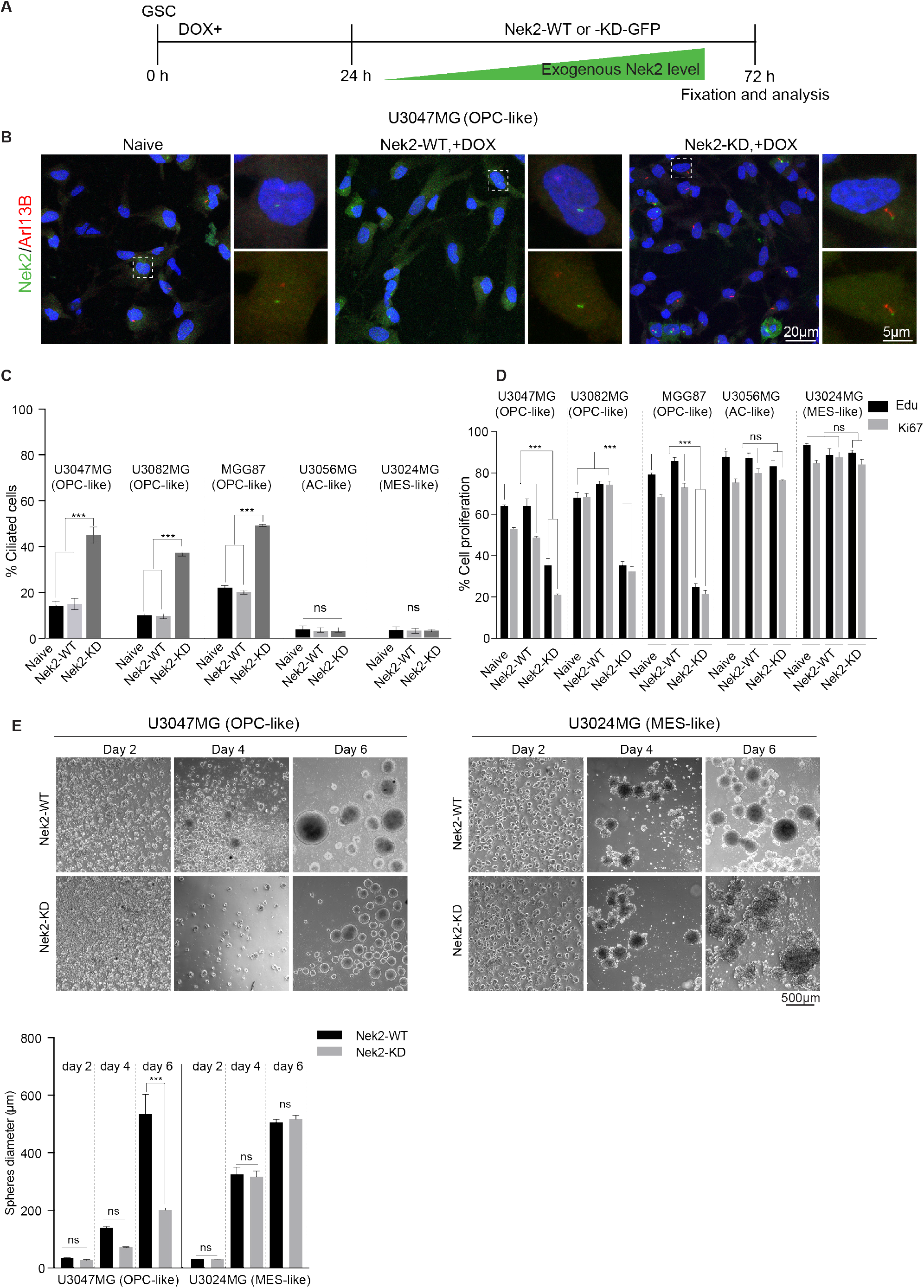
Catalytically inactive Nek2 kinase induces primary cilia in GSCs. **A.** Experimental scheme. Seeded GSCs were induced to express GFP-tagged Nek2-WT and Nek2-KD (kinase-dead, catalytically inactive), fixed, and analyzed. **B.** Compared to naïve or Nek2-WT expressing U3047MG (OPC-like), Nek2-KD expressing cells display cilia. Nek2 (green) labels centrosomes. Arl13b labels cilia (red). Scale bar, 20 μm (overview), 5 μm (Inset). n=4 independent experiments. **C.** The bar diagram quantifies frequencies of ciliated cells across different GSCs upon Nek2-KD expression. Nek2-KD could induce ciliogenesis selectively in OPC-like (U3047MG, U3082MG and MGG87) GSCs and not in MES-like (U3024MG) or AC-like (U3056MG) GSCs. At least 300 cells in each cell line from (n=5) independent experiments. Ordinary two-way ANOVA followed by Tukey’s multiple comparisons test. ***P <0.0001, ns represents non-significant. Error bars show +/− SEM. **D.** Cilium induction impairs the proliferation of OPC-like (U3047MG, U3082MG and MGG87) GSCs but not in MES-like (U3024MG) or AC-like (U3056MG) GSCs. Bar diagram quantifies cell proliferation profile of naïve, Nek2-WT and Nek2-KD expressing GSC lines revealed by 48h EdU pulse chasing Ki67 immunoreactivity. At least 400 cells in each cell line from (n=4) independent experiments. Ordinary two-way ANOVA followed by Sidak’s multiple comparisons test. ***P <0.0001, ns represents non-significant. Error bars show +/− SEM. **E.** Cilium induction impairs neurosphere formation of OPC-like U3047MG but not MES-like U3024MG. Bar diagram quantifies sphere diameter of Nek2-WT and Nek2-KD expressing cells. At least 500 spheres in each cell line from (n=3) independent experiments. Ordinary two-way ANOVA followed by Sidak’s multiple comparisons test. ***P <0.0001, ns represents non-significant. Error bars show +/− SEM.

The finding that responder lines exhibiting reduced proliferation upon cilium induction prompted us to speculate that these GSCs presumably do not form spheres, a typical characteristic of GSCs (Singh et al., 2003; Singh et al., 2004). Our sphere formation assay indicated that U3047MG GSCs formed spheres and progressively grew up to 600 μm in size. In contrast, Nek2-KD expressing GSCs failed to grow beyond 200 μm. Importantly, control experiments that used a non-responder line (U3024MG) that did not respond to cilium induction continue to grow regardless of Nek2-KD expression **(Figure 3Ei-iii).** Since perturbation of CDC induces cilium only in OPC-like GSCs, our further in-depth characterizations have been focused on U3047MG, a representative OPC-like GSC line.

To understand the plausible mechanisms of cilium induction, we analyzed the CDC proteins’ levels before and after cilium induction. Western blots of U3047MG cell extracts revealed an overall reduction in CDC protein levels after cilia induction **(Figure S4A).** We reasoned that this could be due to the general decrease in proliferation and retention of cells at G1-G0, prompting us to estimate the individual CDC component’s recruitment to the basal bodies. Immunostaining using specific antibodies revealed that CDC components’ recruitment is drastically reduced after cilium induction **(Figure S4B)**.

### Newly induced cilia are persistent and structurally and functionally normal

Because cilium induction in responder lines impairs GSCs proliferation **(Figure 3D)**, we conducted a FUCCI-based analysis on U3047MG cells. This analysis revealed that a significant fraction of U3047MG cells reside at G1/G0 after cilia induction **(Figure S5A)**, prompted us to speculate that Nek2-KD-induced cilia are persistent, and thus cells presumably did not proceed to G2-M transition. To test this, we withdrew doxycycline from both responder (U3047MG) and non-responder (U3024MG) GSC lines after day three and continued culturing for a prolonged period of up to 10 days. Although Nek2 expression is unnoticeable at day 10, the abundance of ciliated cells in U3047MG cultures remain unchanged. Concurrently, cell proliferation did not continue. In contrast, the non-responder control U3024MG that could not be induced with cilia continued to proliferate. These findings indicate that newly induced cilia are persistent and suggest that once ciliated, these GSCs do not re-enter the cell cycle **(Figure 4A).**

**Figure 4.**
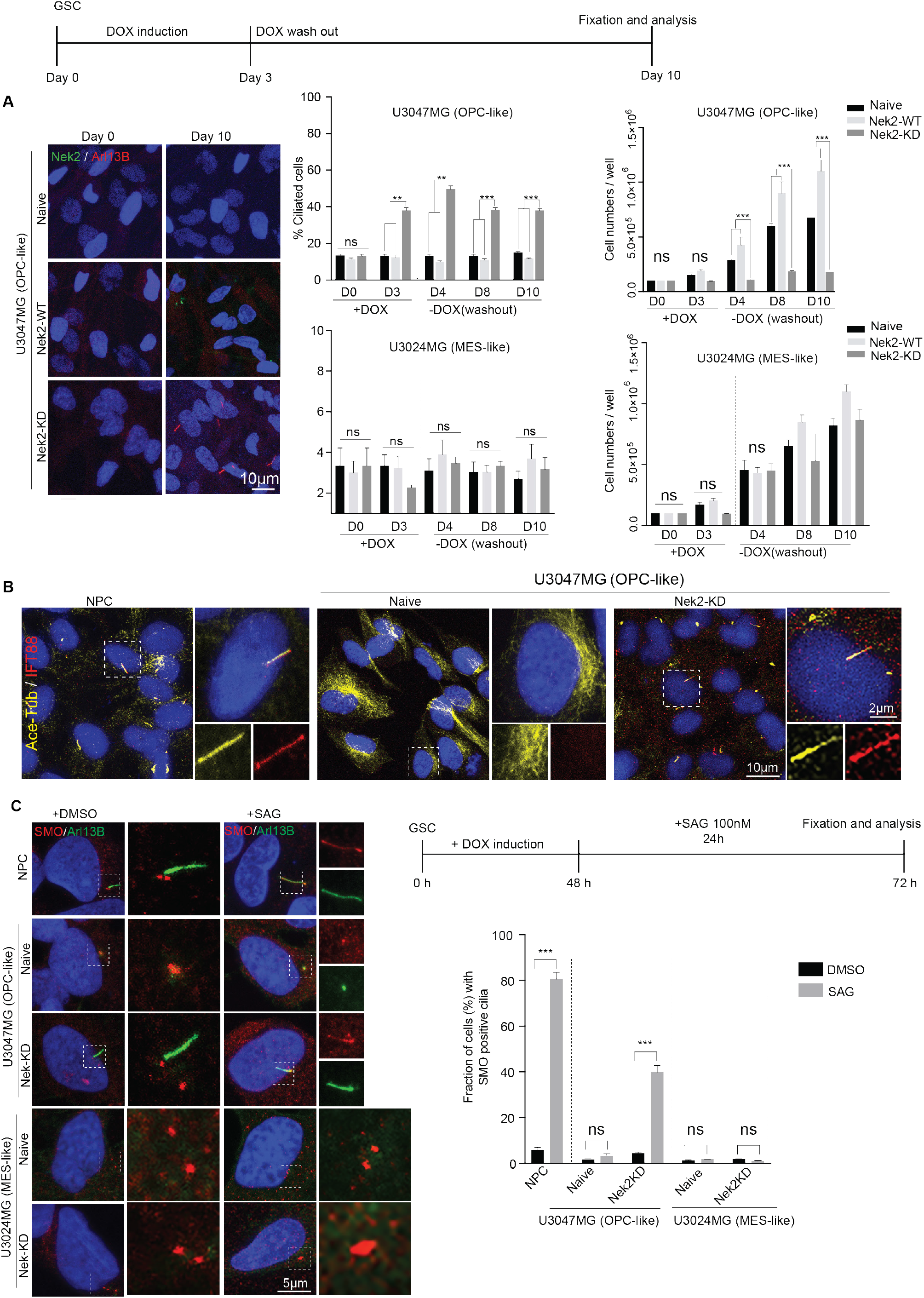
Newly induced cilia in GSCs are persistent and normal in structure and function. **A.** Experimental scheme of doxycycline induction and wash out. In the responder line (OPC-like U3047MG) Nek2-KD expressing cells still harbors primary cilia at day 10 even after removing doxycycline (Scale bar 10 μm). Bar diagram at right quantify the fraction of ciliated cells and absolute cell numbers at various time points of doxycycline wash out. Note that non responder line (MES-like U3024MG) in which cilium could not be induced continue to expand. At least 500 cells in each cell line from (n=3) independent experiments. Ordinary two-way ANOVA followed by Tukey’s multiple comparisons test. ***P <0.0001, **P <0.001, ns represents non-significant. Error bars show +/− SEM. **B.** Just like healthy NPCs (left), newly induced cilia of U3047MG cells (right) display IFT88 particles (red). Cilia are labeled by acetylated α-tubulin (yellow). Naïve U3047MG cells do not display cilia nor IFT88 (middle). Both overview and magnified single cells are shown. Scale bar 10 μm. At least 500 cells from (n=3) independent experiments were examined. **C.** Experimental set up of Shh signaling activation with SAG. In healthy NPCs, SMO (red) is restricted to basal bodies. Upon SAG addition, SMO translocates to primary cilia (top panel). Similar to healthy NPCs, U3047MG (responder) lines induced with cilia but not U3024MG (non-responder) lines display SMO translocation to newly induced cilia (red, bottom panel). Cilia are labeled with Arl13B (green). Naïve U3047MG cells do not show cilia, and thus SMO is restricted to basal bodies (middle panel). Scale bar 5 μm. Bar diagram at right quantifies fractions of cells whose cilia respond to SAG activation. At least 200 cells from (n=3) independent experiments were tested. Ordinary two-way ANOVA followed by Sidak’s multiple comparisons test ***P <0.0001, ns represents non-significant. Error bars show +/− SEM.

Turning our analysis at the ultrastructural level, we analyzed centrioles of U3047MG at an early stage of ciliogenesis where centriole assembles ciliary vesicle and anchors at the membrane serving as a basal body (Moser et al., 2014). Compared to NPCs, centrioles of U3047MG contained swollen and misshaped ciliary vesicles revealing a suppressed ciliogenesis at the early stages of cilium formation. Besides, we also noticed the frequent presence of electron-dense satellite-like particles concentrated at the vicinities of basal bodies, which may be corroborated with an elevated level of CDC components in un-ciliated naïve GSCs **(Figure S5Bi-ii).** On the other hand, we noticed that newly induced cilia structurally appear normal with an infrequent presence of electron-dense particles **(Figure S5iii).**

To analyze the functionality of newly induced cilia, we first tested for the presence of intraflagellar protein-88 (IFT88) and identified that the newly induced cilia were immunoreactive to IFT88 antibodies **(Figure 4B).** Next, we tested whether newly induced cilia can transduce sonic hedgehog (Shh) signaling because it acts through primary cilia (Goetz et al., 2009; Bangs and Anderson, 2017). Smoothened (Smo) is an integral part of the Shh pathway, which relocates to the cilium after activated by a smoothened agonist (SAG). Upon adding SAG, just like healthy NPCs, the activated Smo in U3047MG GSCs relocated from the basal body to the entire length of the cilium in GSCs **(Figure 4C).** These results collectively reveal that perturbing CDC recruitment is sufficient to induce structurally and functionally normal cilia in GSCs.

### Cilium induction triggers GSCs differentiation

To dissect the consequences of cilium induction on GSCs, we tested whether they undergo differentiation. To our surprise, in contrast to unciliated naïve GSCs, GSCs (responder lines of U3047MG, U3082MG, and MGG87) induced with cilia exhibited significantly increased proportions of GFAP, S100β, and TUJ1-positive cells, which are the identity markers of astrocytes and neurons. However, we could not detect any of these markers in the non-responder control U3024MG in which Nek2-KD expression did not induce cilia **(Figure 5A-B).** Importantly, responder GSC lines exhibited a strongly reduced immunoreactivity to Nestin, Pax-6, Sox2, and CD133, which specify neural and cancer stem cells **(Figure 5A-C).** These findings indicate that cilium induction can determine GSCs’ fate and that the differentiation process might be cilia dependent.

**Figure 5.**
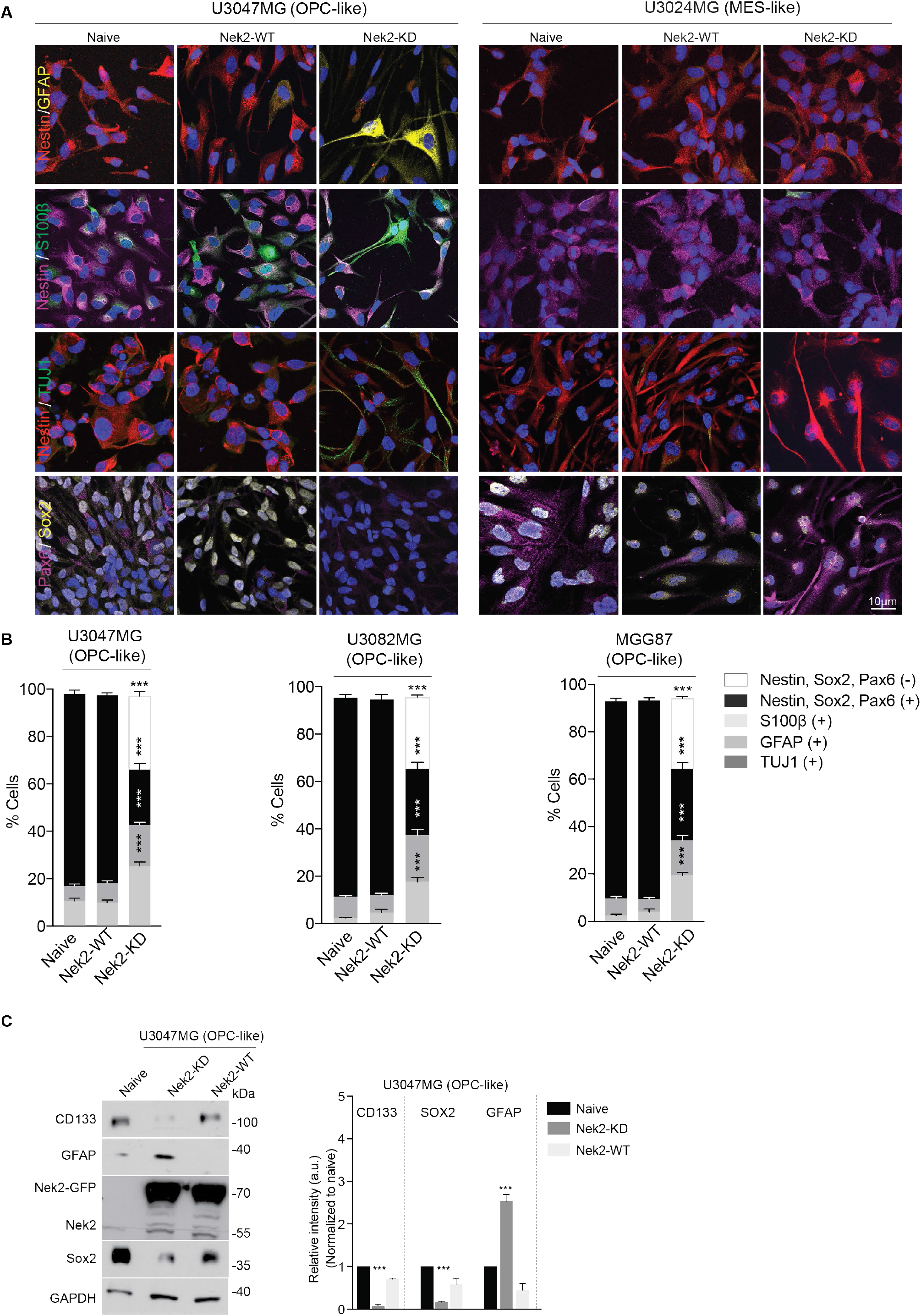
Cilium induction triggers GSCs differentiation. **A.** Naïve or Nek2-WT expressing U3047MG GSCs (responder) display neural stem cell marker Nestin (red), Pax6 (magenta), and Sox2 (yellow) but not differentiation markers of GFAP, S100β, and TUJ-1 (left and middle panel). These GSCs lose neural stem cell markers upon cilium induction but gain differentiation markers of GFAP, S100β, and TUJ1. Non-responder line (U3024MG), in contrast, neither differentiates nor loses its stem cell markers. **B.** Bar diagram below quantifies fractions of OPC-like GSCs (U3047MG, U3082MG and MGG87) positive for stemness (Nestin, Sox2, and Pax6) and differentiation (GFAP and TUJ1) markers. At least 200 cells in each cell lines from (n=3) independent experiments were analyzed. Ordinary two-way ANOVA followed by Tukey’s multiple comparisons test ***P <0.0001. Error bars show +/− SEM. **C.** Western blot compares relative levels of stemness (CD133 and Sox2) and differentiation marker (GFAP) before and after cilium induction. GAPDH was used as a loading control. Bar diagram at right quantifies relative intensities of respective markers before and after cilium induction. Western blots from three (n=3) independent experiments. Ordinary two-way ANOVA followed by Tukey’s multiple comparisons test, ***P<0.0001. Error bars show +/− SEM.

To further demonstrate that differentiation is a cilium dependent process, we ablated cilia by depleting IFT88 (Loskutov et al., 2018). To this end, we first expressed Nek2-KD in U3047MG cells that were depleted of IFT88. As expected, we detected IFT88 immunoreactivity and the appearance of cilia in control siRNA-treated cultures but not in IFT88 siRNA-treated cultures **(Figure 6A).** We then tested whether Nek2-KD could still induce differentiation of U3047MG cultures unable to assemble cilia due to IFT88 siRNA treatment. We first verified that IFT88 depletion did not induce GSCs differentiation. Importantly, Nek2-KD expression induced differentiation of GSCs into GFAP and TUJ1-positive cells only control siRNA-treated cultures but not in IFT88 siRNA-treated cultures **(Figure 6B and the table summarize the findings).** This data indicates that Nek2-KD induced differentiation process is cilia specific.

**Figure 6.**
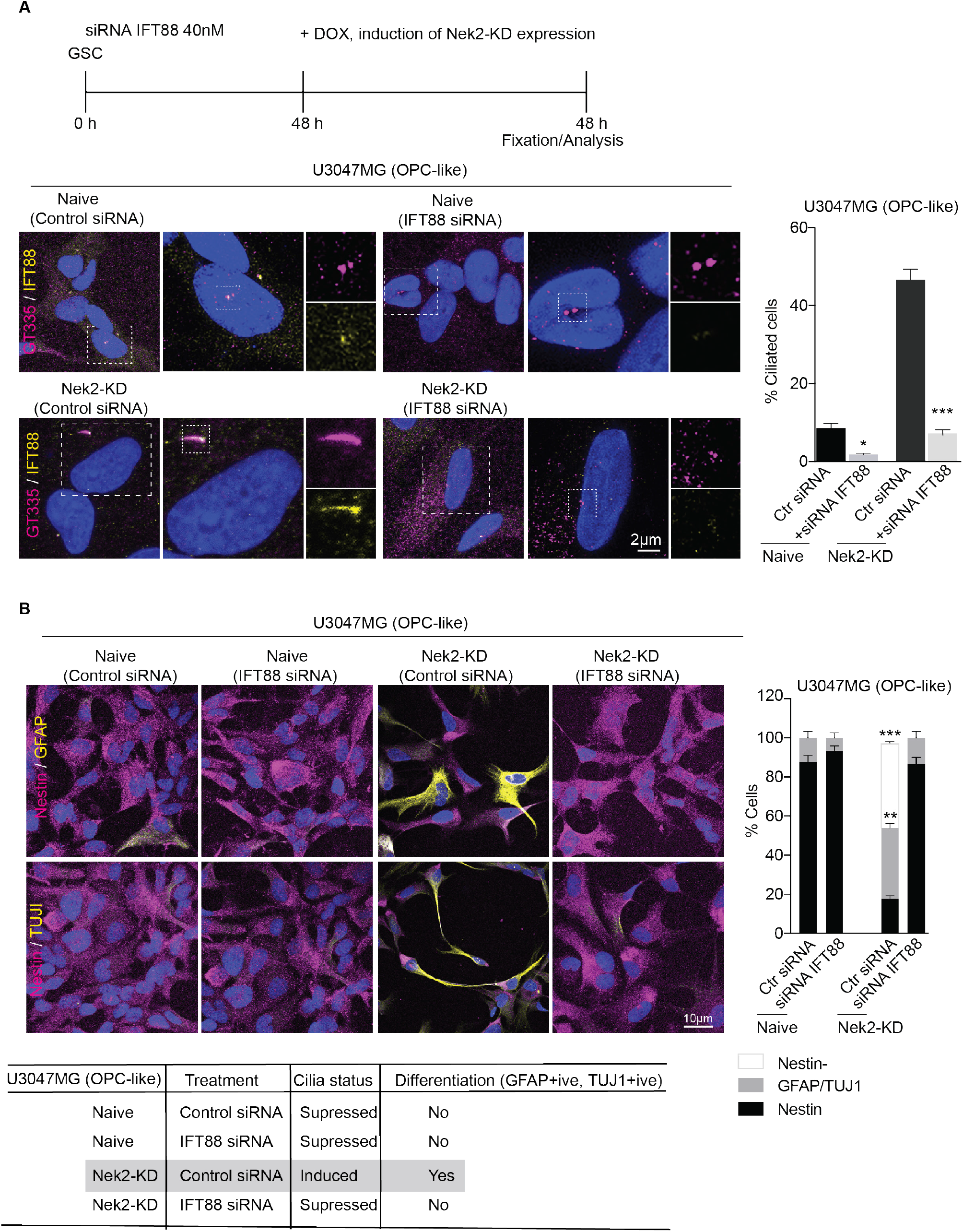
GSCs differentiation is cilium dependent. **A.** Experimental plan. Nek2-KD expression does not induce cilia in GSCs depleted of IFT88. Note Nek2-KD could induce cilia only in control siRNA-treated naïve GSCs cultures in which IFT88 immunoreactivity (yellow) is intact (left panel, both overview and magnified insets are shown). Cilia are labeled with glutamylated tubulin (magenta). In contrast, Nek2-KD does not induce cilia in IFT88 depleted cultures (right panel, both overview and magnified insets are shown). Note that IFT88 expression is lost. Scale bar 2 μm. Bar diagram at right quantifies frequencies of ciliated cells under various conditions tested. At least 300 cells were tested from (n=3) experiments. Ordinary one-way ANOVA followed by Tukey’s multiple comparisons test. ***P <0.0001, *P <0.01. Error bars show +/− SEM. **B.** Nek2-KD expression triggers differentiation (GFAP and TUJ1-positive cells) only in control siRNA treated cells where Nek2-KD could induce cilia (refer to the previous experiment). Thus, Nek2-KD-mediated GSCs differentiation is cilium dependent. Note IFT88 depletion or control siRNA alone does not trigger GSCs differentiation (second and first panel from left). Except Nek2-KD expressing control siRNA-treated cultures, the rest of the cultures exhibit neural stem cell marker Nestin. Scale bar 10 μm. At least 100 cells were tested from (n=3) experiments. Ordinary two-way ANOVA followed by Tukey’s multiple comparisons test. ***P <0.0001, **P <0.001. Error bars show +/− SEM. The table below summarizes the experimental results.

### Cilium induction resets the aberrant PDGFR-α signaling in OPC-like GSCs

To dissect the mechanistic link between cilium induction and GSCs differentiation, first, we investigated the global impact of cilia induction. We analyzed the transcriptome of U3047MG cells before and after cilium induction. We identified >1500 differentially regulated genes **(Figure S6A).** Up-regulated genes were associated with cell morphogenesis in differentiation, whereas down-regulated genes were majorly associated with cell proliferation pathways **(Figure S6B and C).** Likewise, we noticed genes implicated in ciliogenesis were up-regulated, whereas centrosome and CDC component genes involved in cell proliferation and cilia disassembly were down-regulated **(Figure S6D).**

Interestingly, an unbiased analysis of the top 25 differentially expressed genes revealed that SOX10 and PDGFRA are strongly downregulated after cilium induction. Studies have demonstrated that PDGFRA is rapidly downregulated together with oligodendrocyte progenitor (OPC) lineage markers of SOX10 and OLIG2 when OPC differentiate into oligodendrocytes (Dang et al., 2019; Rivers et al., 2008). Notably, down-regulation of stemness maintenance and cell cycle regulators implicate cilium induction switches GSCs from self-renewal to differentiation state **(Figure S6E-F).** Together, the transcriptomic data corroborates with our microscopy-based analysis of cellular identities **(Figure 5A).**

We then analyzed the status of PDGFR-α signaling in U3047MG cells for several reasons. ****i)**** There is a strong association between PDGFR-α overexpression, amplification, and mutation in OPC-like GBMs (Neftel et al., 2019; Verhaak et al., 2010) ****ii)**** We detected an increased level of total PDGFR-α in OPC-like GSCs **(Figure S6F)** and indeed, PDGFR-α signaling is positively correlated to the self-renewal property of GSCs, accelerated tumor onset, and increased tumor invasion (Brennan et al., 2013; Clarke and Dirks, 2003; Filbin et al., 2018; Pathania et al., 2017; Verhaak et al., 2010) ****iii)**** PDGFR-α signaling is regulated by primary cilia in healthy cells (Schmid et al., 2018). ****iv)**** Cilium induction was significant in OPC-like GSCs **(Figure 3C).** Western blot analysis revealed that in contrast to un-ciliated U3047MG cells, ciliated cells showed a drastically reduced level of total PDGFR-α, which in turn correlated to reduced levels of its downstream signaling components such as activated pAKT and c-myc. This finding suggests that newly induced cilia in U3047MG can play a role in sequestering excessive PDGFR-α **(Figure S6G).** Supportive of this is our super-resolution imaging assay strikingly indicating the presence of PDGFR-α, mostly in newly induced cilium. Importantly, naïve U3047MG cells that have suppressed ciliogenesis exhibited unspecific localization of PDGFR-α nearly the entire cell’s space suggesting that cilium induction switches PDGFR-α to be relocated to a newly induced cilium **(Figure S6H).**

From these findings, we reasoned that the up-regulated total PDGFR-α in GSCs is constitutively active in ligand independent manner, which could enable them to continuously self-renew. To test this aspect, we imaged the dynamic localization of PDGFR-α at various time points upon its activation by its endogenous ligand PDGF-AA **(Figure S7).** In control experiments that used healthy NPCs, we detected the appearance of PDGFR-α signal in cilia 20 mins after the ligand addition (Top panel). In contrast, un-ciliated U3047MG cells always displayed excessive PDGFR-α immunoreactivity in a ligand-independent manner (Middle panel). Strikingly, after cilium induction, U3047MG cells showed PDGFR-α dynamics similar to healthy NPCs such that PDGFR-α is directed to a newly induced cilium (Bottom panel). In summary, these results elucidate the dominant effect of cilium induction in triggering differentiation of GSCs via regulating abnormally functioning PDGFR-α signaling.

### Cilium induction impairs GSCs invasion into 3D human brain organoids and mouse brain

To analyze the impact of cilium induction in GSCs invasion in 3D, we adapted our recently optimized brain organoid-based GSCs invasion assays, which recapitulate some of the *in vivo* behavior of GSCs (Goranci-Buzhala et al., 2020; Mariappan et al., 2020; Linkous et al., 2019). In our assays, we seeded 1000 naïve and Nek2-KD expressing U3047MG cells at the vicinity of 10-day-old organoids, differentiated from transgenic iPSCs expressing Tubulin-GFP. For visualization purposes, GSCs were tagged with mCherry reporter **(Figure 7A).** After 24-hrs of seeding, we added doxycycline to induce Nek2-KD protein expression and imaged the invasion behavior of GSCs at various time points from day 1 to 7 in live. Imaging at brightfield, we noticed that on day 2, GSCs were attracted to brain organoids. On day 7, a significant proportion of naïve GSCs have invaded the organoids. Strikingly, Nek2-KD expressing GSCs failed to enter brain organoids. In contrast, the non-responder control U3024MG in which Nek2-KD expression did not induce cilia invaded the organoids **(Figure 7B)**.

**Figure 7.**
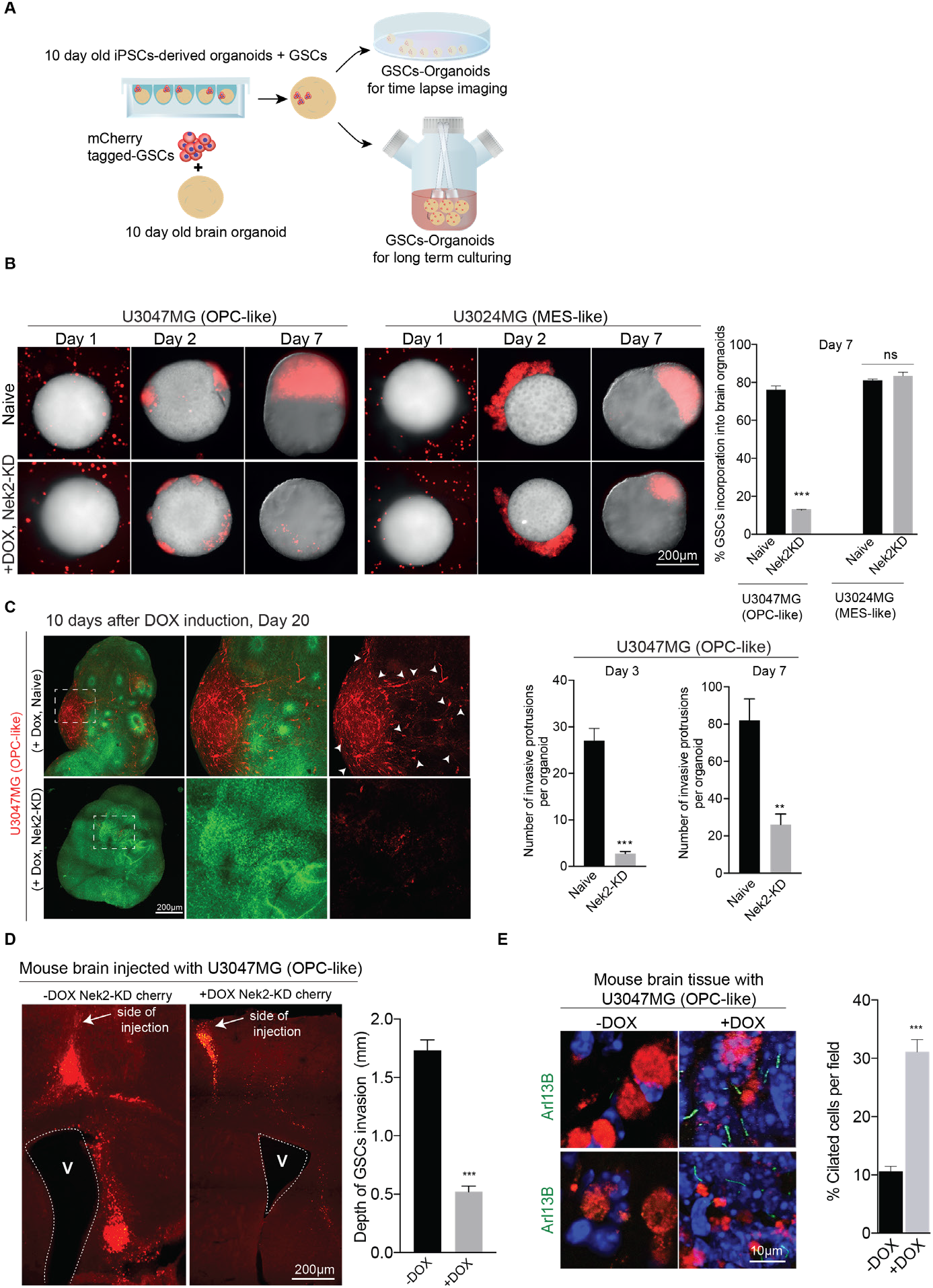
Cilium induction prevents GSCs invasion into 3D human brain organoids and mouse brain. **A.** Experimental scheme showing the process of GSCs invasion assay. Co-culturing of labeled GSCs with 10-day-old brain organoids until imaging them. **B.** Bright-field images show that mCherry tagged OPC-like U3047MG and MES-like U3024MG cells (top panel) invasion into 3D human brain organoids. Upon Nek2-KD induction, OPC-like U3047MG cells (bottom left) exhibit impaired invasion but not MES-like U3024MG cells (bottom right). In these experiments, untagged iPSCs were used to generate brain organoids. Scale bar 200 μm. At least 20 organoids from 5 different batches were tested. Ordinary two-way ANOVA followed by Tukey’s multiple comparisons test ***P <0.0001, ns represents non-significant. Error bars show +/− SEM. **C.** Tissue cleared 3D imaging of the whole organoid after 20 days of GSCs invasion. Tissue clearing enhances the visualization of U3047MG-mCherry growth (red) into the organoids (green). Invading GSCs form protrusions and microtube networks (top panel). In contrast, Nek2-KD expressing U3047MG cells poorly grow within organoids, as evidenced by poor growth and less intensity (bottom panel). The bar graph at right quantifies the parameters of invasiveness before and after cilium induction. Unpaired t-test, ***P <0.0001, **P <0.001. Error bars show +/− SEM. **D.** Mouse brains at 4 weeks after grafting m-Cherry tagged U3047MG cells expressing Nek2-KD. Coronal sections of the grafted brains are shown. The control animal groups that do not drink doxycycline (−DOX) water exhibit invading GSCs deep into the striatum. In contrast, coronal sections of the animal groups that drunk doxycycline (+DOX) exhibit an impaired invasion or migration of GSCs. The arrow marks the site of GSCs injection. Dotted lines mark ventricles (V). Scale bar 200 μm. At right, bar diagram quantifies depth of GSCs invasion between control (that did not drink doxycline, −DOX) and experimental (that did drink doxycycline, +DOX) animal groups. At least ten sections / animal were analyzed from n=4 (control group) and n=5 (experimental group) animals. Unpaired t test. ***P <0.0001. Error bars show +/− SEM. **E.** Immunofluorescence analysis of cilia. GSCs at the control group (that did not drink doxycline, −DOX) do not show cilia. In contrast, GSCs at the experimental animal groups (that did drink doxycycline, +DOX), show newly induced cilia (green). At least ten sections / animal were analyzed from n=4 (control group) and n=5 (experimental group) animals. Scale bar 10 μm. Unpaired t test. ***P <0.0001. Error bars show +/− SEM.

To enhance the image quality and quantitatively determine the 3D invasion behavior of GSCs at the later time point of day-20, we utilized our recently described rapid organoid clearing and imaging methods (Goranci-Buzhala et al., 2020). Naïve U3047MG cells extensively diffused and recapitulated the known characteristics of invading GSCs, such as establishing protrusion-like processes in the form of microtubes (Osswald et al., 2015). In contrast, Nek2-KD expressing U3047MG cells exhibited an impaired organoid invasion and, therefore, failed to grow in brain organoids **(Figure 7C and Movies 1 and 2).**

To determine the fate of Nek2-KD expressing U3047MG cells in 3D organoids, we performed immunofluorescence imaging of thin-sectioned organoids harboring GSCs invasion. We noticed that Nek2-KD expressing U3047MG cells, but not naïve cells were strongly positive for GFAP exhibiting characteristic astrocyte-like cell shapes. These findings reveal that Nek2-KD expressing U3047MG cells underwent a differentiation process similar to what was observed in our 2D experiments upon cilium induction **(Figure S8).** Together, our organoid based invasion assay elucidates that Nek2-KD expression triggers differentiation of GSCs within brain organoids thereby impairs their invasion. To further test if cilium induction can perturb GSCs invasion *in vivo*, we grafted 2×104 Nek2-KD expressing U3047MG cells intracerebrally into immune-deficient mice. This procedure generates infiltrative tumor xenografts closely mimicking the parental tumor’s behavior (see method for details) (Ricci-Vitiani et al., 2010). To induce cilia via Nek2-KD expression, we supplied animals 1mg/ml doxycycline in drinking water. At week four after grafting, control groups that did not drink doxycycline water displayed GSCs along the needle tract spreading onto the corpus callosum and deeper into the striatum. Conversely, experimental groups that drunk doxycycline water harbored only a few GSCs at the grafting area. Most of them were embedded in cell debris and did not spread further to the corpus callosum and striatum **(Figure 7E).** Examining them for cilium induction revealed that U3047MG cells displayed cilia, which was abnormally long, corroborating that cilium induction can significantly perturb U3047MG invasion in *vivo* **(Figure 7F).**

## Discussion

Glioblastoma constitutes self-renewing, highly tumorigenic GSCs, which exhibit striking similarities to NPCs expressing neural stem cell markers, assemble spheres, and rapidly self-renew (Bhaduri et al., 2020; Rajakulendran et al., 2019). Besides, patient-derived GSCs rapidly invade into human brain organoids and mouse brain, assemble invasive networks, and form tumors that phenocopy patient tumors (Goranci-Buzhala et al., 2020; Mariappan et al., 2020; Linkous et al., 2019; Osswald et al., 2015) **(Figure 7).** Identifying a unique feature of GSCs that strikingly differ from NPCs and exploiting the molecular regulation may provide hints to the development of strategies to selectively impair GSCs proliferation and invasion.

NPCs are functionally characterized by their abilities to self-renew and differentiate. Recently, others and we have identified that the primary cilia are a molecular switch whose spatiotemporal dynamics decisively regulate self-renewal versus differentiation of NPCs (Gabriel et al., 2016; Kim et al., 2011; Li et al., 2011). Delayed cilia disassembly triggers NPCs differentiation, whereas accelerated cilia disassembly promotes NPCs proliferation (Gabriel et al., 2016; Kim et al., 2011; Li et al., 2011). This provides conceptually a novel ‘cilium checkpoint’ that can be targeted to regulate stem cell fate. Although GSCs are strikingly similar to NPCs, GSCs possibly differ in at least two aspects. First, cilium formation is suppressed at the early stages of ciliogenesis due to elevated levels of molecular players (CDC components) that trigger cilia disassembly **(Figure 1 and 2).** Second, GSCs are characterized by continuous self-renewal and block differentiation (Park et al., 2017; Rajakulendran et al., 2019).

We reasoned that suppressed ciliogenesis might provide a selective advantage to GSCs to continuously self-renew, evading the cell cycle exit and differentiation in a cilium checkpoint independent manner. It is important to emphasize that cilia loss has a severe consequence in triggering melanoma metastasis by promoting WNT/beta-catenin signaling and resistance development (Zhao et al., 2017; Zingg et al., 2018). Thus, the primary cilium could function as a tumor suppressor organelle in broader cancer types. However, to date, no effort has been made to test the hypothesis of whether reintroducing cilia can be anti-tumorigenic, triggering differentiation of stem cells and impair their invasion. Therefore, we targeted the cilia disassembly mechanisms of GSCs **(Hypothesis Figure S1).** Our work identifies that depleting CDC levels as a mechanism to induce cilia in GSCs persistently. Inducing ciliogenesis in a subset of patient-derived GSCs has triggered them to lose their stemness and gain differentiation programs **(Figure 5 and S6).** As a result, GSCs induced with cilia failed to invade into 3D human brain organoids and mouse brains **(Figure 7).**

Recruitment of CDC components to the ciliary base is associated with temporal cilia disassembly at the onset of cell cycle re-entry (Gabriel et al., 2016). However, the molecular interplay between CDC recruitment and cilia disassembly remained elusive. It is conceivable that the CDC could contain an enzyme that can be activated to depolymerize the microtubule cytoskeleton of cilia during disassembly. The identification of Kif24, a depolymerizing microtubule kinesin, and Nek2 as components of CDC helps to unveil the mechanisms of CDC-mediated cilia disassembly (Kim et al., 2015). It is noteworthy that Kif24 and its physiological activator Nek2 kinase levels are elevated in naïve GSCs that exhibit suppressed ciliogenesis **(Figure 2).** The depletion of cell cycle-related kinase has also been shown to induce cilia in a small fraction of serum-starved U251MG cells (Yang et al., 2013). It is noteworthy that, unlike GSCs, the commercially available U251MG cells were cultured in the presence of serum. Thus, it remains unclear whether the small fraction of ciliated cells observed is due to the depletion of kinase or serum starvation.

Inducing ciliogenesis, GSCs behave like NPCs undergoing a differentiation suggesting that newly induced cilia are structurally and functionally normal. Proving this is the newly induced cilia harbor IFT88 and transduces ligand-dependent activation of Shh-signaling **(Figure 4).** This supports the notion that cilia can functionally be restored in a subset of GSCs that are categorized as OPC-like GSCs expressing an elevated level of PDGFR-α. Restoring cilia first sequesters excessive PDGFR-α into newly induced cilia and resets aberrant PDGFR-α signaling **(Figure S6G-H).** The observed drastic reduction in total PDGFR-α levels after cilium induction is a surprising phenomenon suggesting that cilia could harbor proteasomal subunit components to degrade newly translocated ciliary PDGFR-α. Interestingly, at least three components of 19S proteasomal subunits have been identified in mouse embryonic fibroblast cilia (Gerhardt et al., 2015). It is thus tempting to investigate if cilia of NPCs and GSCs harbor proteasomal subunits and E3 ligases that can recognize PDGFR-α and critically regulate its amount and downstream signaling.

PDFGRA signaling is mediated by the primary cilium (Schneider et al., 2005). Of note, expressing Nek2-KD sensitized only PDGFR-α overexpressing OPC-like GSCs but not other subtypes. There could be several reasons for this selectivity. Compared to other CDC components, OPC-like GSCs (responder lines) express the highest level of Nek2 **(Figure 2A-B).** Second, it is possible that OPC-like GSCs (which exhibit an increased level of PDGFRA) still harbor cilium induction programs that are perhaps masked in other cellular states. Finally, PDGFR-α overexpressing GSCs may harbor epigenetically regulated neurogenic and astrogenic features. Thus, restoring ciliogenesis could favor differentiation and loss of self-renewal. For example, the transcription factor ASCL1 in OPC-like (proneural subtypes) GSCs has been shown to unlock chromatin allowing new sites to activate differentiation programs (Park et al., 2017). Interestingly, we noticed that the responder line (U3047MG) do express quantitatively a higher ASCL1 expression than the non-responder line (U3024MG) (Data not shown). Recent work has also uncovered an elevated level of EZH2, a Polycomb Repressive Complex 2, as a mechanism by which cancer cells lose cilia and promote metastatic melanoma (Zingg et al., 2018). Future experiments with detailed transcriptomic and systems analysis will uncover yet unknown epigenetic factors that mask cilium induction and differentiation in a wide variety of GSCs.

Currently, it remains unknown why cilium induction was not possible in non-responder of AC-like and MES-like lines. The non-responder lines may likely harbor defective centrioles, which will not assemble cilia at all. Thus, a comprehensive analysis of centriole, basal body, and cilia structure is required. Second, it is unknown whether mutations in cilia inducing genes and if they are under epigenetic regulation. Finally, it remains unknown if ciliary microtubule polymerases (such as Kif24) are constitutively active in non-responder lines.

In summary, as a cellular organelle that regulates signaling, self-renewal, and differentiation, the cilium checkpoint emerges as an attractive molecular switch that can be targeted to alter the fundamental aspects of GSCs. Identification of CDC components and their elevated levels coupled with suppressed ciliogenesis begins to explain why GSCs tend to continuously self-renew and invade into the brain. Cilium-induced differentiation in GSCs has a broader impact on 3D brain organoids, which has recently received considerable attention as a human in vitro system to evaluate GSCs invasion and glioma-neuron interaction (Goranci-Buzhala et al., 2020; Linkous et al., 2019). As shown in our organoid-based invasion assays and mouse xenografts, invasive protrusions formed by naïve GSCs were completely abrogated upon cilium induction **(Figure 7)**. Thus, perturbed GSCs invasion observed in neural epithelial tissues of brain organoids and mouse brain suggests that targeting CDC, cilium checkpoint including enzymes that deconstruct cilia, emerge as “Achilles heel” to limit or prevent GSCs invasion.

## Supporting information

Movie 1 Naive

Movie 1 Nek2-KD

RNA seq table

## Acknowledgement

We thank the members of the Laboratory for Centrosome and Cytoskeleton Biology and the Institute of Human Genetics for their support. We thank Dr. Hiroaki Wakimoto for offering GSCs (MGG87, MGG8). We extend our gratitude to Prof. Dieter Häussinger and Dr. Boris Görg for providing their support with their microscope facility. We thank Abida Pranty Islam for helping with biochemical experiments. JG and GG are supported by a DFG grant (GZ: GO 2301/5-1). RP is supported by AIRC grant (IG 2019 Id.23154). GG is supported by the Pharmacology and Bayer Graduate Program at the University of Cologne.

## Author contribution

JG, AM, and GG conceived the concept of project. GG performed most of the experiments. JG supervised the work. AM performed super-resolution, organoid imaging and shRNA mediated CDC depletion. GC and MG performed electron microscopy analyses. AP and NJ performed RNA transcriptomic analyses. JG wrote the manuscript and inputs were given by all the authors.

## Supplementary figures and legends

**Figure S1.**
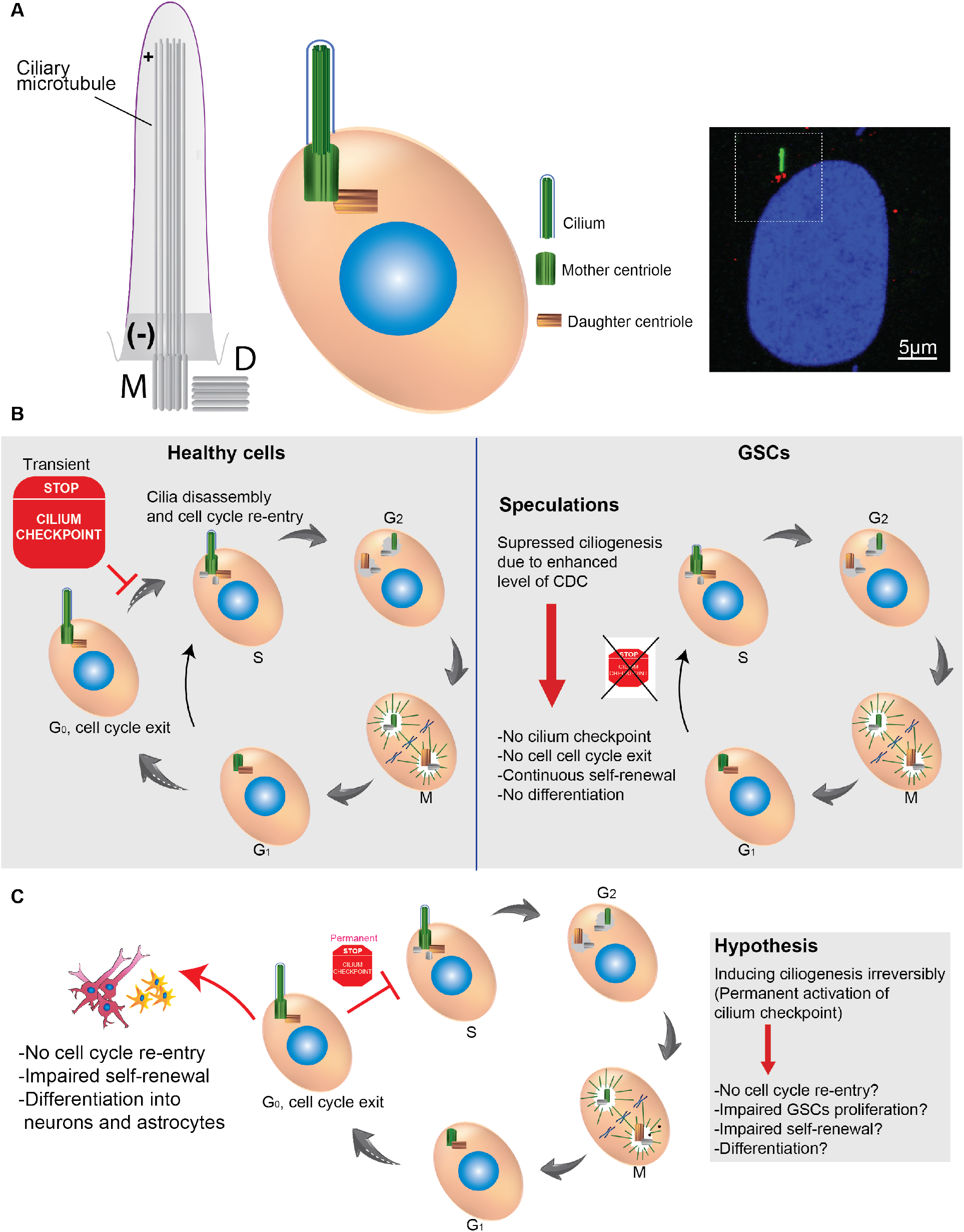
Cilium checkpoint and hypothesis. **A.** The cartoon depicts the structure of primary cilium and a cell with primary cilium. Mother centriole (basal body) templates the formation of the ciliary microtubule. Both the positive and minus end of the ciliary microtubule is labeled. Legends are given at right. A representative image of a cell with primary cilium (Arl13, green) and basal body (γ-tubulin, red) is given at right. **B.** Cilium checkpoint in healthy cells (left panel). Cilium is dynamic. The mature mother centriole serves as a template for cilium formation at the onset of cell cycle exit (G0). The cilium begins to disassemble at the onset of cell cycle re-entry (S). Failure or delay in cilium disassembly acts as a transient brake “cilium checkpoint” in cell cycle progression (stop sign). Cilium disassembly at G2 triggers cells to continue with mitotic progression. In GSCs (Speculations, right panel) ciliogenesis is suppressed due to CDC elevation. Therefore, GSCs do not have cilium checkpoint, and this could result in no cell cycle exit, continuous self-renewal, and no differentiation. **C.** Working hypothesis. Inducing cilia could permanently activate cilium checkpoint. As a result, GSCs may not re-enter the cell cycle, exhibit impaired proliferation, self-renewal, and undergo terminal differentiation into astrocytes and neurons.

**Figure S2.**
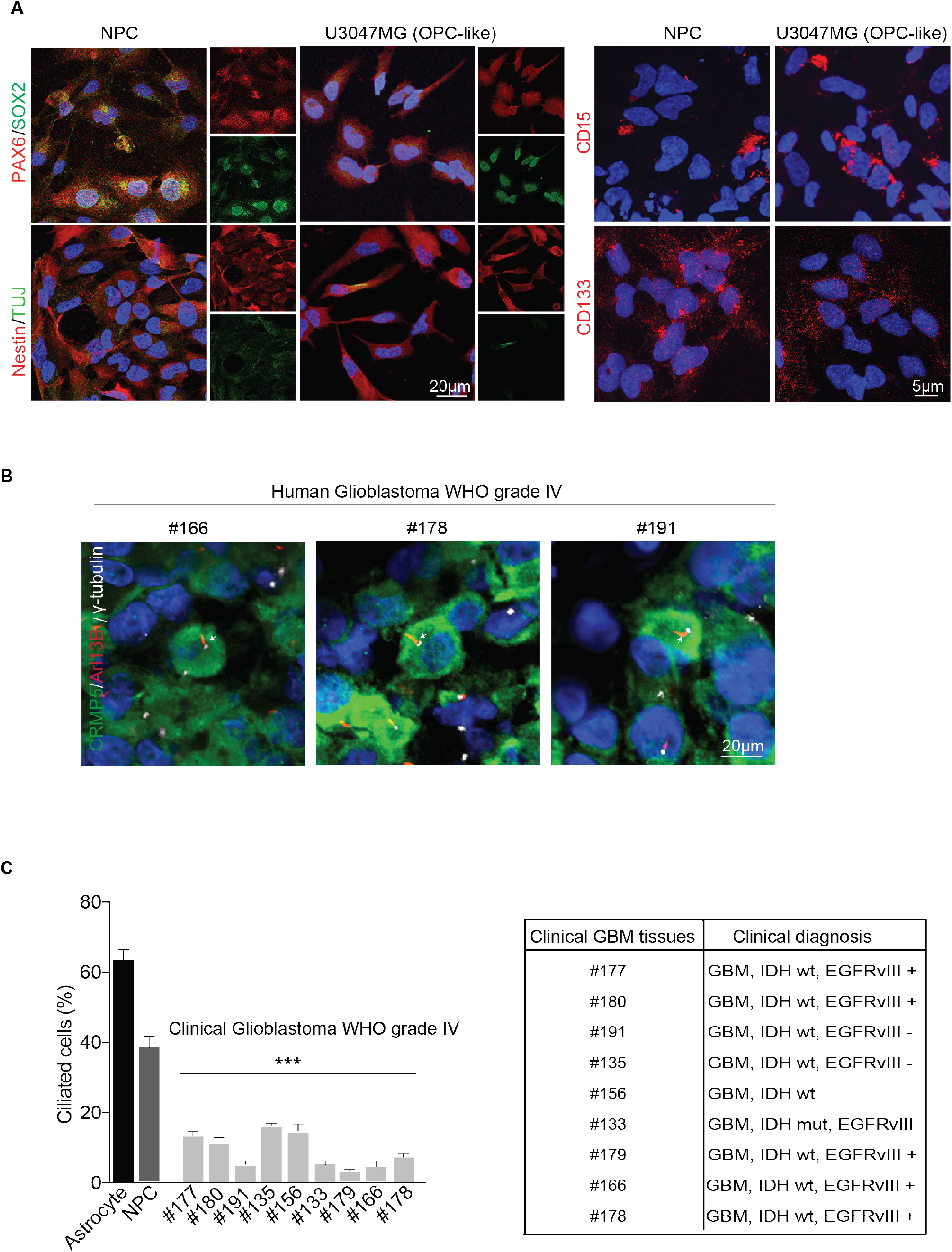
Patient-derived GSCs have neural stem cell attributes and exhibit suppressed ciliogenesis. **A.** Similar to NPCs, patient-derived GSCs express neural stem cell markers such as Pax6, Sox2, Nestin, including CD15 and CD133. U3047MG is used as representative GSCs. Representative images from at least 4 independent experiments. Scale bar, 20 μm (overview), 5 μm (high magnification). **B.** Related to main figure 1B. Cilia occurrence in clinical tissues. Grade IV glioblastoma tissues mostly lack cilia (red, arrows), γ -tubulin labels centrosomes (basal bodies, white). CRMP5 marks cancer cells in clinical tissues (green). **C.** The bar diagram quantifies frequencies of ciliated cells. Note that in clinical tissues, at least 9 independent glioblastoma grade IV tissues show an overall reduction in ciliation compare to astrocytes and NPCs. Tumor cells were scored based on CRMP5 immunolabeling. At least 500 cells in each cell line from three (n=3) independent experiments. One-way ANOVA followed by Dunnett’s multiple comparisons test ***P <0.0001. Error bars show +/− SEM. Table on the right shows clinical information on tissue samples.

**Figure S3.**
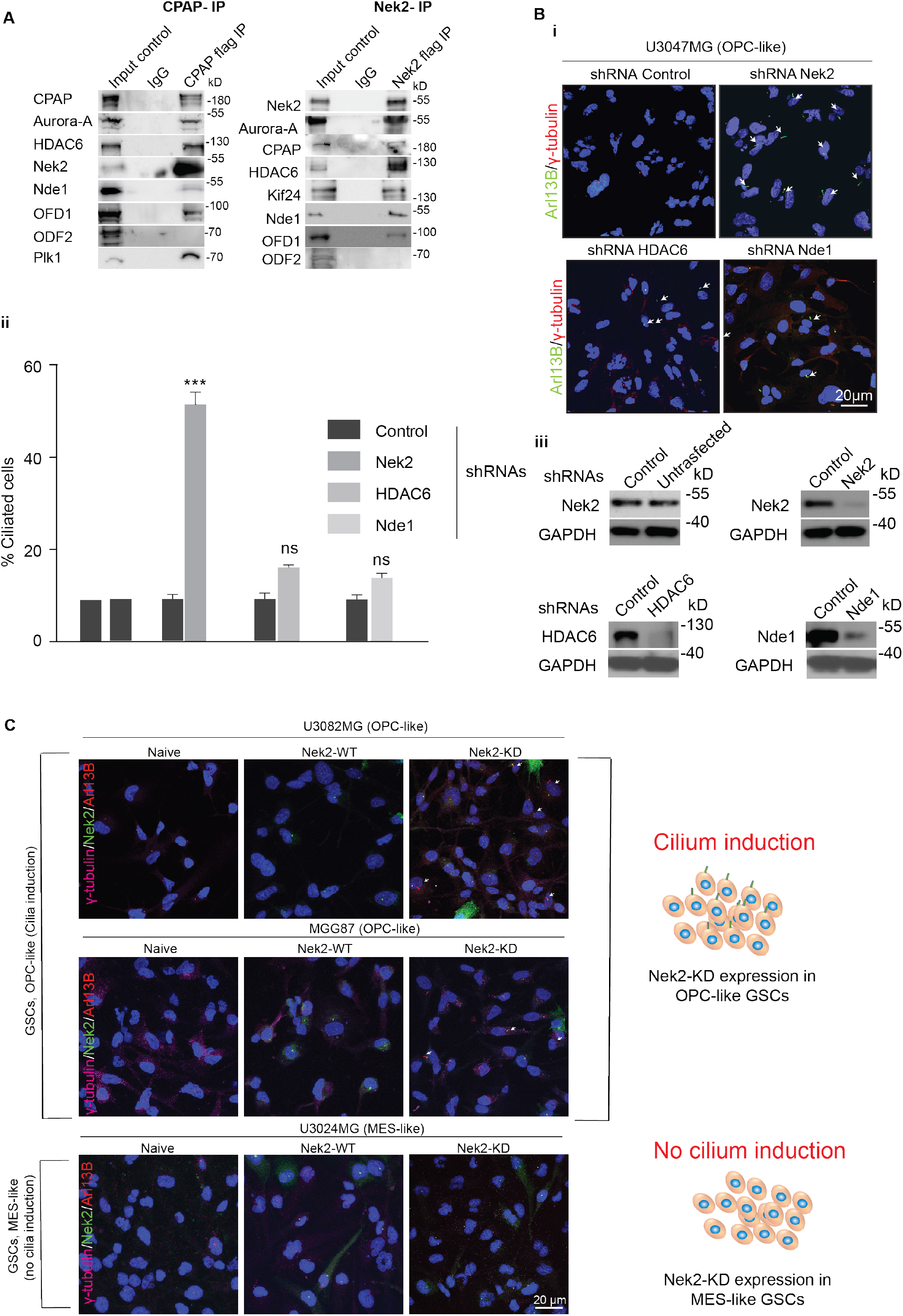
Depletion of CDC components induce cilia in patient-derived GSCs. **A.** Right panel. Immunopurification of CPAP complex from HEK293 cell extracts expressing CPAP-flag. Nek2 co-purifies with other components of CDC. Centrosome-associated Outer Dense Fiber protein-2 (OFD-2) does not co-purify with CPAP indicating the specificity of CPAP interaction with CDC components, including Nek2. IgG indicates negative control. Left panel. Reciprocal immunopurification experiment. Immunopurification of Nek2 complex from HEK293 cell extracts expressing Nek2-flag. Nek2 co-purifies with other components of CDC, including Kif24. IgG indicates negative control. **B. i.** shRNA specifically targeting Nek2, HDAC6, and Nde1 depletes respective proteins and induce cilia in U3047MG cells. U3047MG is used as representative GSCs. Arl13B is applied to label cilia (green) and γ-tubulin is used to label the centrosomes (basal bodies, red). Scale bar, 10 μm. **ii.** Bar diagram quantifies frequencies of ciliated cells in cultures treated with shRNA specific to CDC components. **iii.** Western blots below show shRNA-mediated depletion of specific CDC components. GAPDH is a loading control. At least 200 cells were tested from (n=3) experiments. Ordinary two-way ANOVA followed by Sidak’s multiple comparisons test. ***P <0.0001, ns represents non-significant. Error bars show +/− SEM. **C.** Related to main Figure 3C. Catalytically inactive Nek2 kinase (Nek2-KD) could induce cilia in OPC-like GSCs but not in MES-like GSCs (U3024MG). Arl13B labels cilia (red, white arrows), Nek2-WT, and KD both basal target bodies (green), and γ-tubulin labels the centrosomes (basal bodies, magenta). Scale bar, 20 μm.

**Figure S4.**
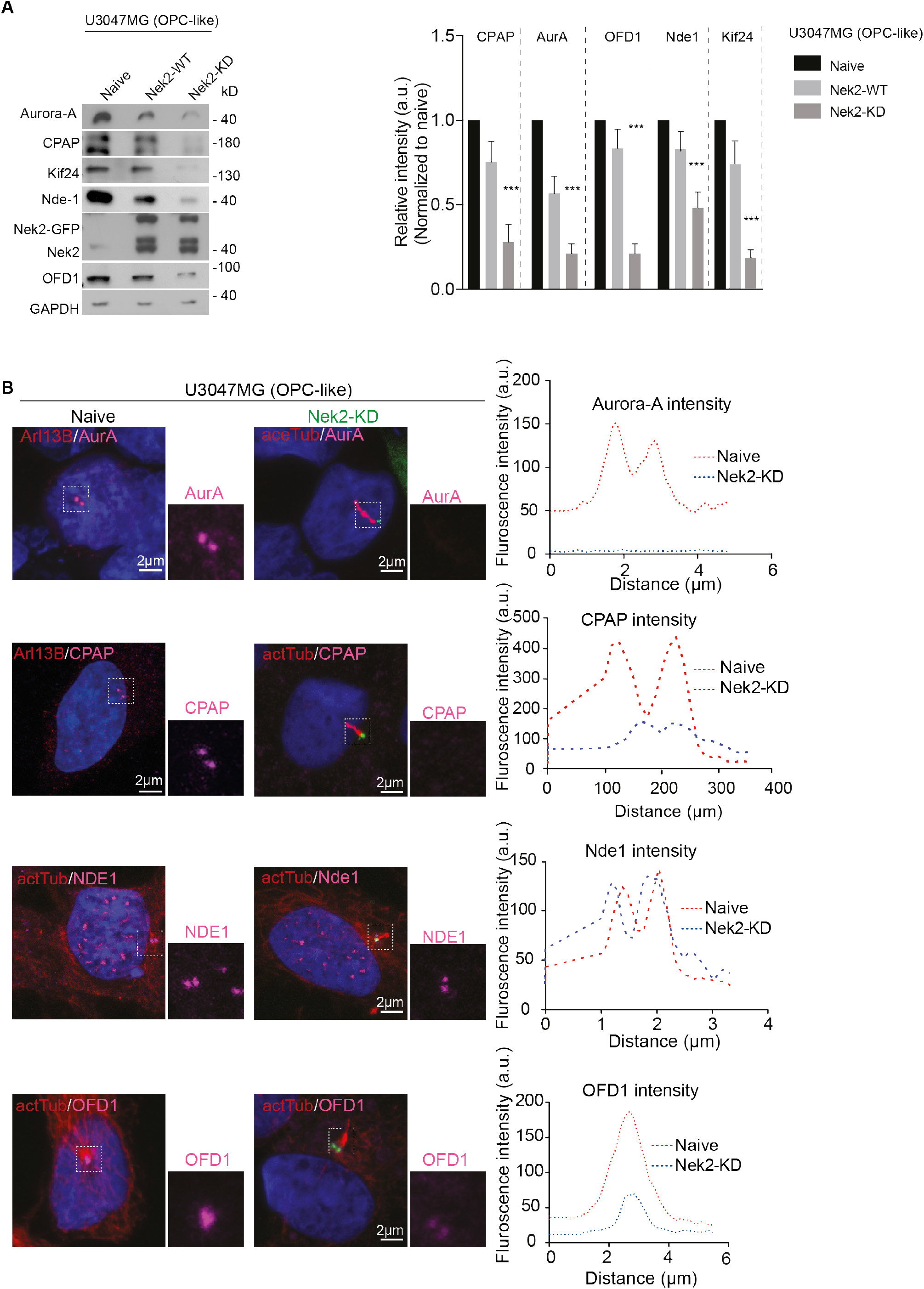
Nek2-KD expression impairs CDC recruitment to induce primary cilia. **A.** Semi-quantitative Western blot exhibiting relative levels of CDC components before and after cilium induction in U3047MG cells. Cilium induction (Nek2-KD expressing U3047MG) is associated with a reduction in overall CDC component levels. Nek2 antibody recognizes both endogenous and exogenous proteins. GAPDH was used as a loading control. The bar diagram at right quantifies the relative intensity of CDC components before and after cilium induction. Western blots from three (n=3) independent experiments. Ordinary two-way ANOVA followed by Tukey’s multiple comparisons test, ***P<0.0001. Error bars show +/− SEM. **B.** Compared to non-ciliated U3047MG naïve, Nek2-KD expressing ciliated U3047MG cells recruits a reduced level of CDC components. Cilia are labeled either by Arl13B or acetylated α-tubulin (red). CDC components (magenta) were immunostained using antibodies specific to Aurora-A, CPAP, Nde1, and OFD1. Nek2-KD labels centrosomes at the base of cilia (green). Graphs at right display the distribution of fluorescence intensity of CDC components recruited to ciliary base before (red, naïve) and after (blue, Nek2-KD) cilium induction. At least 300 cells in each cell line from (n=4) independent experiments. Scale bar 2 μm.

**Figure S5.**
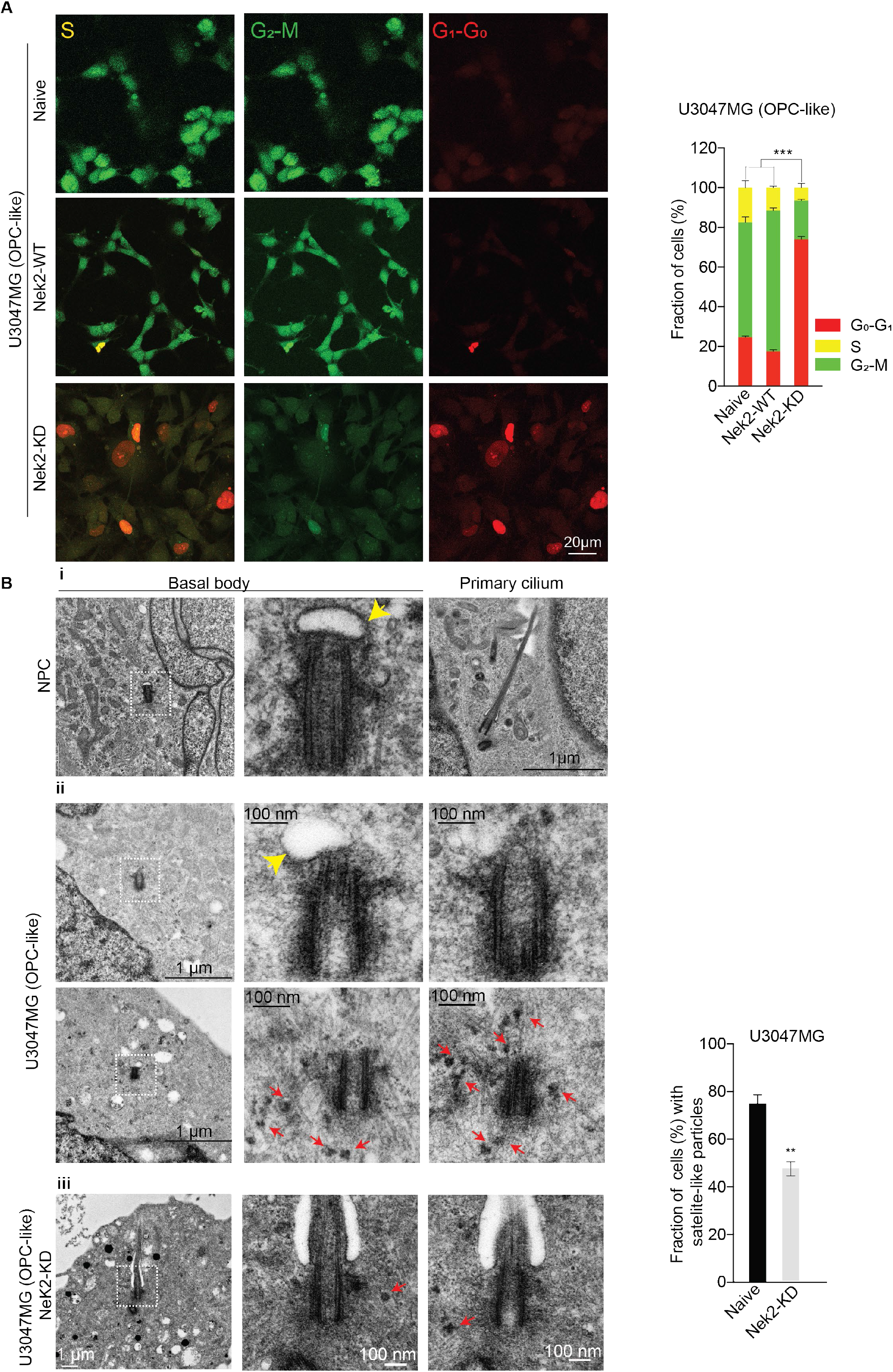
Cilium induction impairs GSCs proliferation and retains the majority of cells at the cell cycle exit stage of G0-G1. **A.** Most non-ciliated naïve or Nek2-WT expressing U3047MG cells reside at G2-M phase of the cell cycle (proliferating stage, green). In contrast, most ciliated Nek2-KD expressing U3047MG cells reside at G1-G0 phase of the cell cycle (cell cycle exit stage, red). The graph at right segregates proportions of GSCs retained at different stages of the cell cycle. At least 100 cells were tested from (n=3) experiments. Ordinary two-way ANOVA followed by Tukey’s multiple comparisons test. ***P <0.001. Error bars show +/− SEM. **B.** Ultra-structural analysis of healthy NPCs **(i)** non-ciliated naïve U3047MG **(ii)** compared to ciliated U3047MG cells **(iii)**. Misshaped vesicles (yellow arrow) and numerous electron-dense satellite-like particles are prevalent (red arrow) in naïve U3047MG that are infrequently seen after cilium induction and in healthy NPCs. The bar diagram at right quantifies it. At least 20 centrioles were analyzed. Scale bar 1 μm (low magnification) and 100nm (high magnification). Unpaired t-test. **P<0.001. Error bars show +/− SEM.

**Figure S6.**
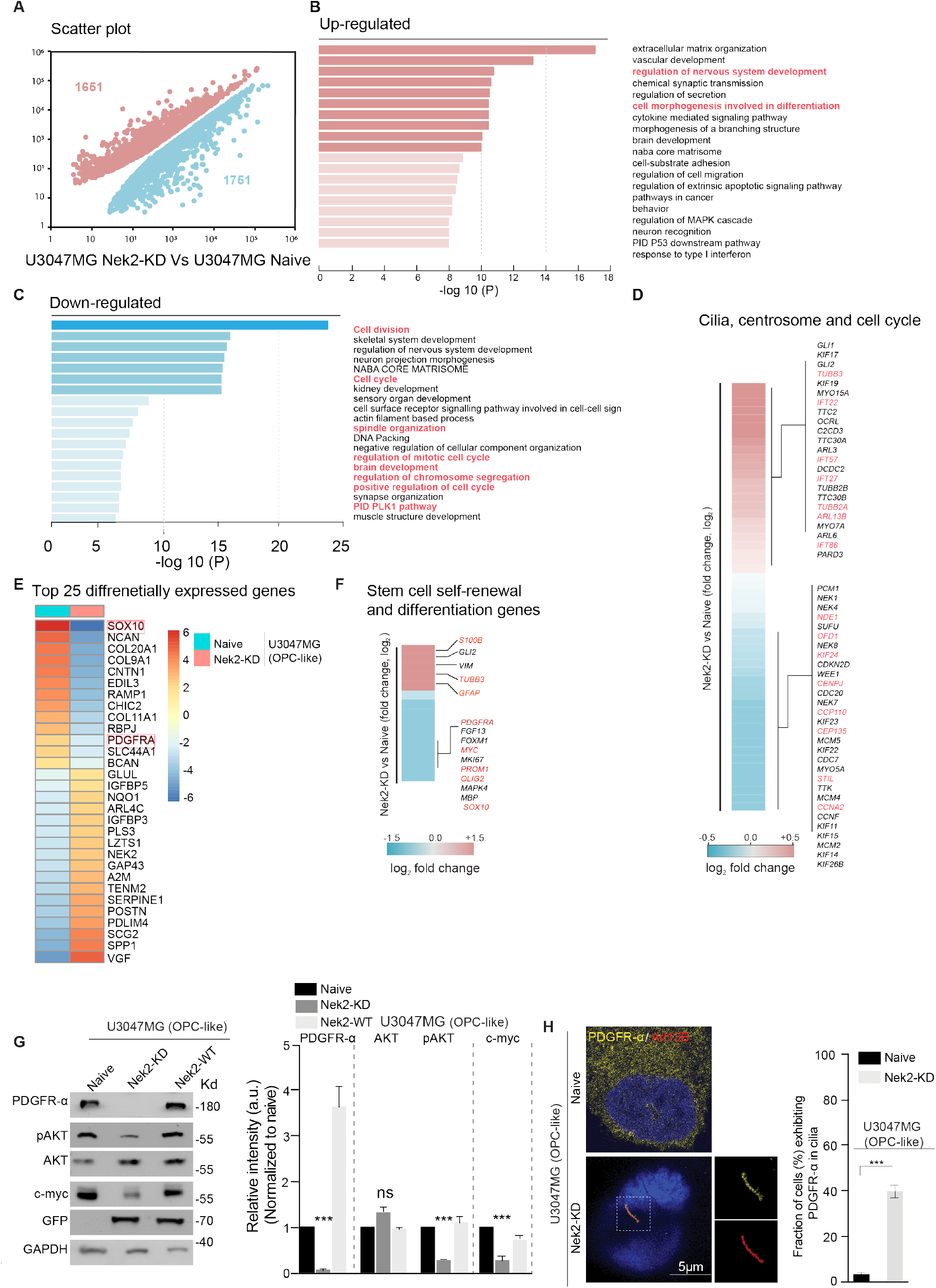
Transcriptomic analysis before and after cilium induction indicates the signs of cilium-dependent GSCs differentiation. **A.** Scatter plot of changing gene expression levels (FPKM; >0.6 log2-fold change, FDR<0.01) in U3047MG naïve), ciliated GSCs (U3047MG NEK2-KD) cells. **B.** The top most enriched GO terms associated with the 1651 upregulated genes in Nek2-KD expressing U3047MG cells. Critical pathways related to cell cycle progression and differentiations are marked in red fonts. **C.** As in panel B, but for the 1751 downregulated genes in U3047MG Nek2-KD cells. Critical pathways related to cell cycle progression and differentiations are marked in red fonts. **D.** Log2-fold gene expression changes of genes associated with cilia, centrosome, and cell cycle regulation according to GSEA classification. **E.** Unbiased top 25 genes differentially expressed before (naïve) and after cilium induction (Nek2-KD) shows SOX10 and PDGFRA are among the down regulated after cilium induction **F.** Genes associated with. Selected genes related to stem cell, self-renewal and cell differentiation are marked in red fonts. Fold-change ±0.6 log2 adjusted P-value of <0.05. **G.** Western blot compares relative levels total PDGFR-α and its downstream signaling effectors in GSCs before and after cilium induction. Note PDGFR-α level is drastically reduced after cilium induction (Nek2-KD expressing GSCs). Similarly, active AKT and c-myc levels are also reduced. GAPDH was used as a loading control. Bar diagram at right quantifies relative levels of PDGFR-α and its downstream signaling effectors before and after cilium induction. Western blots from three (n=3) independent experiments. Ordinary two-way ANOVA followed by Sidak’s multiple comparisons test, ***P<0.0001, ns represents non-significant. Error bars show +/− SEM. **H.** Newly induced cilium sequesters PDGFR-α. Non-ciliated naïve U3047MG displays an all over distribution of PDGFR-α (yellow, left panel). Upon cilium induction, excessive PDGFR-α is directed to the newly induced cilium. Unpaired t test. ***P <0.0001. Error bars show +/− SEM. Scale bar 5 μm.

**Figure S7.**
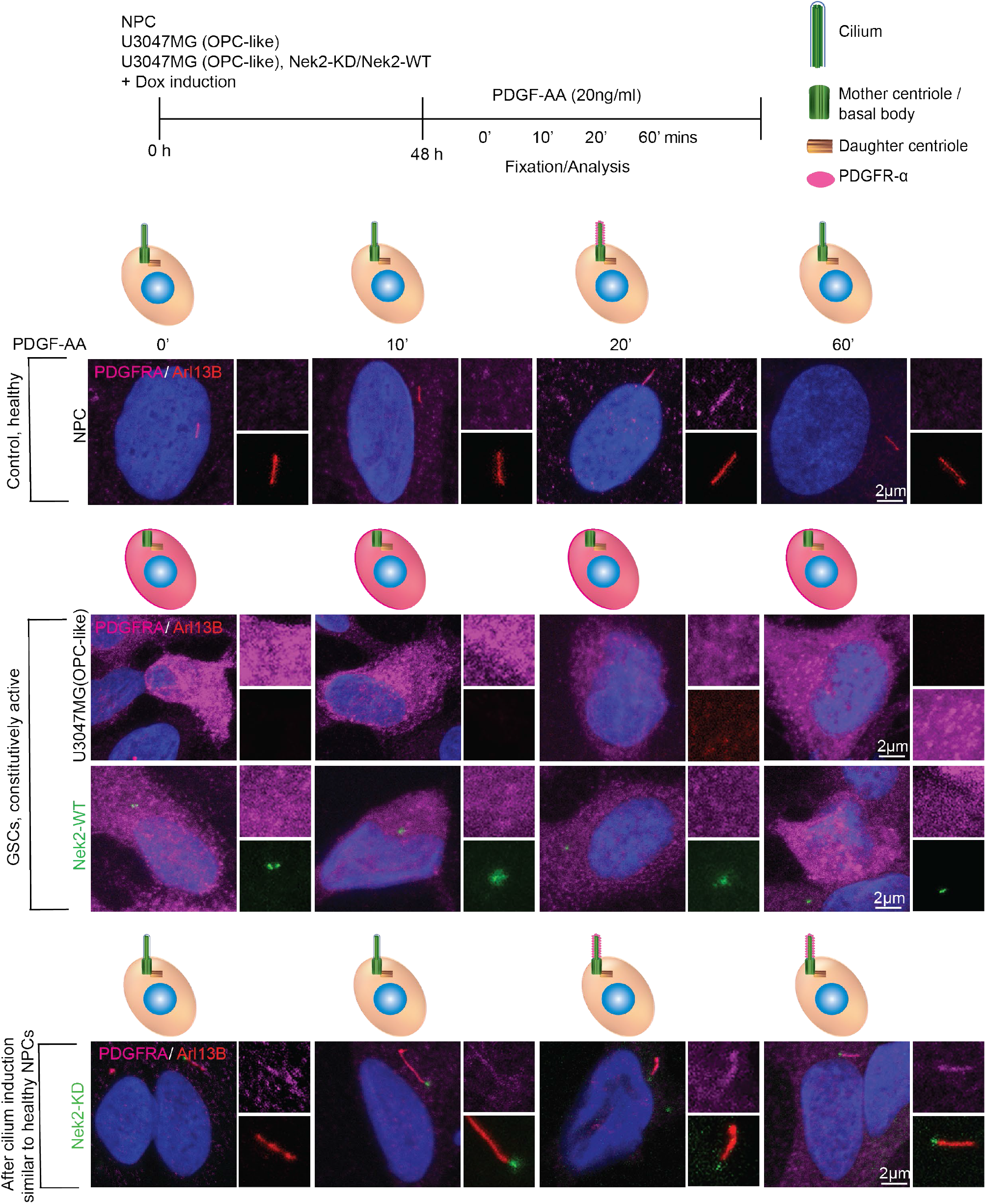
PDGFR-α in OPC-like GSCs is constitutively active. Top panel. Experimental plan. In healthy NPCs, PDGFR-α is activated within 20 mins of PDGF-AA addition. Note that activated receptor localizes at the primary cilium. Arl13B labels cilia (red) and PDGFR-α is stained using specific antibodies (magenta). The cartoon is given to simplify the visualization of PDGFR-α dynamics. Scale bar 2 μm. At least 200 cells were tested from (n=3) experiments. Middle panel. Naïve or Nek2-WT (Green) expressing U3047MG (OPC-like) cells exhibit strong PDGFR-α immunoreactivity (magenta) regardless of PDGF-AA addition, indicating that PDGFR-α in these cell types is constitutively active in a ligand-independent manner. Nek2-WT (green) labels centrosomes. The cartoon shows PDGFR-α distribution. Scale bar 2 μm. At least 100 cells were tested from (n=3) experiments. Bottom panel. Nek2-KD (green) expressing ciliated U3047MG (OPC-like) cells behaves just like healthy NPCs and exhibits similar PDGFR-α dynamics upon ligand addition. A visibly apparent level of PDGFR-α is directed to newly induced cilium at 60 mins of PDGF-AA addition. Arl13b labels cilia (red) and PDGFR-α is stained using specific antibodies (magenta). Nek2-WT (green) labels basal bodies. Cartoon simplifies the visualization of PDGFR-α dynamics. Scale bar 2 μm. At least 200 cells were tested from (n=3) experiments.

**Figure S8.**
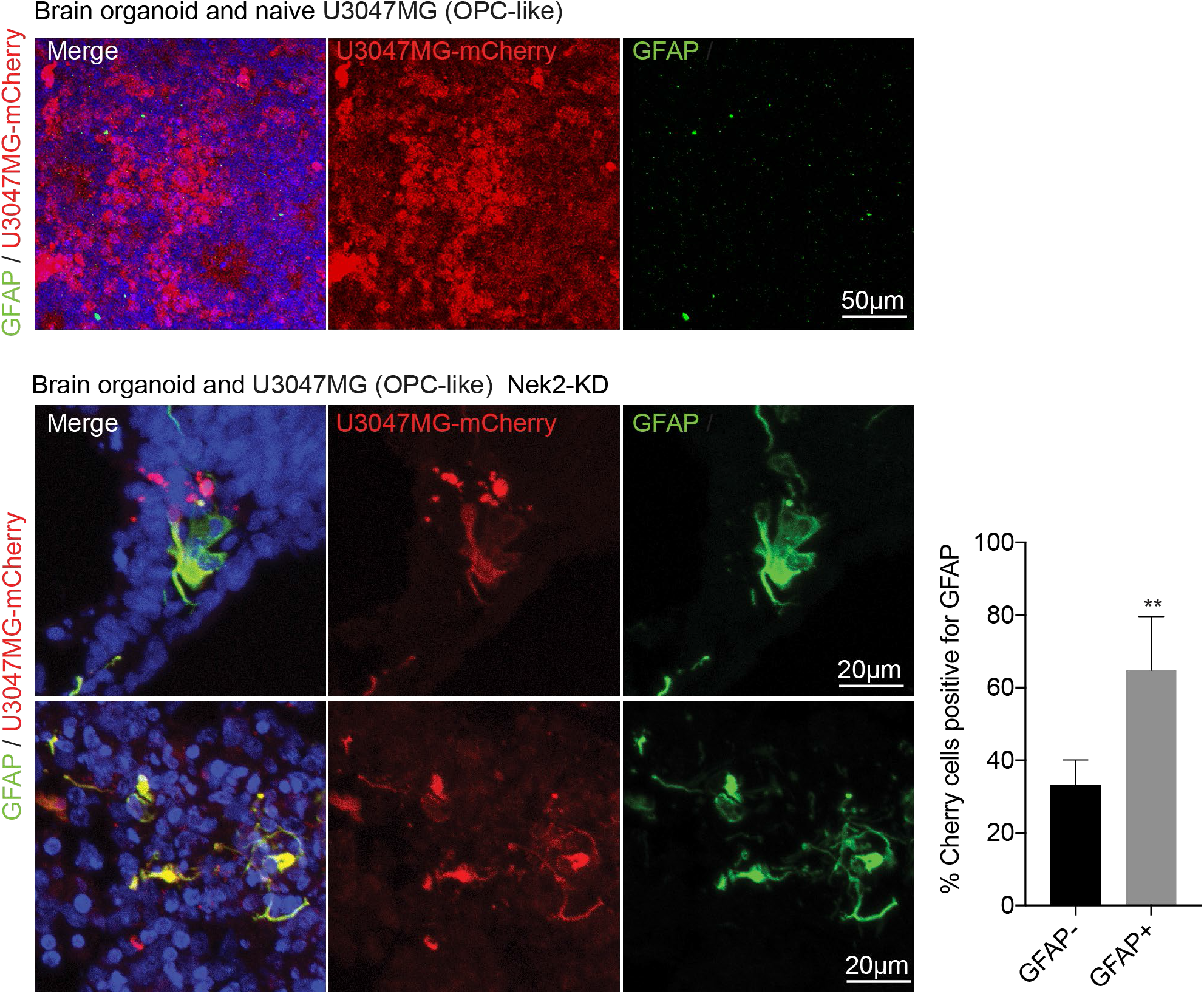
Cilium induction prevents GSCs invasion into 3D human brain organoids. Immunostaining of thin-sectioned organoid slices exhibits diffused growth of U3047MG-mCherry (top panel). Note, U3047MG-mCherry cells do not differentiate to GFAP-positive cells at this condition. Scale bar 50 μm. U3047MG-mCherry (red) cells undergo differentiation to GFAP-positive cells (green) upon Nek2-KD induction. Note U3047MG-mCherry cells at this condition do not proliferate. Instead, they undergo differentiation as judged by their co-localization with GFAP (green). Graph at right quantifies GFAP-negative and -positive cells. Scale bar 20 μm. At least 100 cells were tested from 5 independent organoids. Unpaired t-test, **P <0.001. Error bars show +/− SEM.

**Movie 1.** Related to main Figure 7B. Representative image of tissue cleared 3D imaging of whole organoid (GFP) after 20 days of U3047MG naïve (mCherry) invasion. *Z-*stacks movie show typical ventricular zones of brain organoid, extensive microtube networks and invasive protrusions of naïve U3047MG cells. Scale bar, 100 μm

**Movie 2.** Related to main Figure 7B. Representative image of tissue cleared 3D imaging of whole organoid (GFP) after 20 days of U3047MG Nek2-KD (mCherry) invasion. *Z*-stacks movie show Nek2-KD expressing U3047MG cells (mCherry) exhibited an impaired invasion and failed to grow in brain organoids (green). Scale bar, 100 μm

**Table 1. RNA seq**

## Methods

### Plasmids and cloning

For lentiviral transductions, full-length Nek2-WT and Nek2-KD (K32R) were subcloned from pLVX 3xflag plasmid into a lentiviral packing vectors pLenti6.3 and pLIX_403. These constructs contain the hPGK promoter and with C-terminal GFP. The pLVX 3xflag NEK2-WT and pLVX 3xflag NEK2-KD was generously provided the Daynlacht lab (Kim et al., 2015). For immunoprecipitation experiments, GFP-tagged CPAP was subcloned from pSIN CPAP-WT plasmid into a lentiviral vector pSIN containing the CMV promoter and in-frame N-terminal 3xflag tag.

### Primary GSCs isolation from GBMs

Primary GSCs from the GBMs were isolated using an established protocol (D’Alessandris et al., 2017; Ricci-Vitiani et al., 2010). In brief, GBM surgical specimens were mechanically dissociated and cultured in DMEM /F12 (Gibco) medium containing 2 mM glutamine, 0.6% glucose, 9.6 g/mL putrescine, 6.3 ng/mL progesterone, 5.2 ng/mL sodium selenite, 0.025 mg/mL insulin, and 0.1 mg/mL transferrin sodium salt (Sigma-Aldrich), supplemented with 10ng/ml epidermal growth factor (EGF) (Peprotech) and 10ng/ml basic fibroblast growth factor (bFGF) (Peprotech). Typically, GSCs are characterized by forming spheres expressing stem-cell markers, such as CD133, sex-determining region Y-box 2 (Sox2), Musashi, and Nestin. CD133 expression was detected by anti-CD133–phycoerythrin (AC133-PE) antibody or PE-conjugated mouse immunoglobulin (Ig)G1 isotype control antibody (Miltenyi Biotec). The expression of Sox2 was analyzed by PerCP-Cy 5.5 mouse anti-Sox2 or PerCP-Cy 5.5 mouse IgG1 isotype control (Becton Dickinson [BD]). Viable cells were identified using 7-amino actinomycin D (Sigma-Aldrich). Cells were analyzed with a FACSCanto flow cytometer (BD).

Besides sphere formation assay, stemness phenotypes of GSCs were assessed by self-renewal capacity and generation of xenografts that are histologically mimicking the parent tumor. To assess clonogenicity, viable cells were dispensed at different densities in 96-well plates by fluorescence-activated cell sorting (FACS Aria, BD). When required, subtype classification of GSCs was performed with the RT2 Profiler PCR Arrays (Qiagen) (Marziali et al., 2017). All of the described experiments have been performed low passaged GSC cultures.

### Culturing and maintenance of patient-derived GSCs

As previously described, patient-derived GSCs (used in Figure. 1) were cultured in Neurocult NS-A basal medium (STEMCELL technologies) supplemented with 10% BSA (STEMCELL technologies), L-Glutamin (Gibco, 25030081), Heparin (STEMCELL technologies), B27 without vitamin A (Gibco), N2 (Gibco), 20ng/ml recombinant human epidermal growth factor (hEGF) (Peprotech), 20ng/ml basic fibroblast growth factor (bFGF) (Peprotech) (Xie et al, 2015) (Marziali et al., 2017). Cells were maintained at 37°C and 5% CO2 on poly-L-ornithine-coated (Sigma) and laminin-coated (Sigma) cell culture dishes. Cells were dissociated using Accutase (Thermo Fisher). GSCs were cultured in stem cell medium as described above and analyzed for expression of stem cell markers such as, SOX2, PAX6, Nestin, and for the absence of differentiation markers such as TUJ1 and GFAP.

In ligand stimulation experiments, cells were stimulated with 20 ng/ml PDGF-AA ligand (R&D Systems) for 10–60 min. To study Shh signaling, the agonist SAG (Tocris) 100nM was used. After treatments, cells were fixed and analyzed for respective markers. To analyze cell proliferation, EdU was pulsed in cells for 24h before fixation and stained according to the manufacturer’s instructions (Click-iT EdU assay kit, Life Technologies).

### Primary GSCs and GBM tissues used

At least eight patient-derived GSC lines were used in this study. U3047MG (PDGFRA, OPC-like), U3082MG (PDGFRA, OPC-like), MGG87 (PDGFRA, OPC-like), #450 ((PDGFRA, OPC-like), UG3056MG (EGFR, AC-like), U3024MG (NF1, MES-like), #275 (CDK4, NPC-like) (Xie et al., 2015) (Neftel et al., 2019). At least nine GBM grade IV tissue sections were used. GBM grade IV, #177 (IDH wt, EGFRvIII+), #180 (IDH wt, EGFRvIII+), #191 (IDH wt, EGFRvIII-), #135 (IDH wt, EGFRvIII-), #156 (IDH wt), #133 (IDH mut, EGFRvIII-), #179 (IDH wt, EGFRvIII-),#166 (IDH wt, EGFRvIII+), #178 (IDH wt, EGFRvIII+).

### Real-time quantitative PCR

Total RNA was extracted from the patient derived GSCs using RNeasy mini extraction kit (Qiagen Cat No./ID: 74104) according to the manufacture’s instruction. Equal amounts of the different samples of amplified RNA (1000 ng) were transcribed into cDNA. The reverse transcription (RT) reaction was carried out using SuperScript™ VILO™ cDNA Synthesis Kit (Invitrogen). Gene expression was measured by performing TaqMan PCR using Gene Expression MasterMix (Applied Biosystems) for PDGFRA (Hs00998018_m1), EGFR (Hs01076090_m1), CDK4 (Hs00364847_m1), NF1 (Hs01035108_m1) on the 7500 Real-Time PCR System (Applied Biosystems) supplied with 7500 Real-Time software. Gene expression fold changes were calculated using the ΔΔC_T_ method and GAPDH (Hs 02758991_g1) was used as a housekeeping gene.

### Lentiviral production and transduction of target cells

Constitutive overexpression of mCherry U3047MG lentiviral vectors was prepared using pSicoR-Ef1a-mCh-Puro (Addgene plasmid 31845). For GSCs, stable lines generation inducible lentivectors were used pLix 603 Nek2-KD EGFP and pLix 603 Nek2-KD EGFP.

The lentiviral shRNA constructs pLKO.1-puro for CDC components (NDE1, OFD1, HDAC, Nek2) were kindly provided by Dr. Rajalingam. The cloned vectors were packed into lentivirus using second-generation packaging plasmids (pMD2.G, Addgene #12259 and psPAX2, Addgene #12260). Briefly, target vectors and packaging plasmids were transfected into HEK293TS cells using calcium chloride. After 16 h, the medium was changed, and the virus was collected after 8 h. The freshly harvested virus was used to transduce target cells in a 1:1 ratio for 72–96 h.

To generate GSCs lines stably expressing inducible GFP tagged-Nek2-WT and Nek2-KD, pLIX_403 lenti inducible gateway cloning vector (Addgene #41395) was used. The target cells were transduced with lentivirus containing pLIX-NEK2-WT and pLIX-NEK2-KD and selected with puromycin antibiotic 2μg/ml. For the expression of the transgenes, all the cell lines stably expressing inducible lenti vector were induced with 2-5 μg/ml of doxycycline (Sigma) for 24–96 h.

### Culturing and maintenance of human iPSC culture

At least three different human iPSCs were used in this study, namely, mEGFP (AICS-0012), mRFP (AICS-0031) TUBA1B, and IMR90 (Wicell). Cells were plated on Matrigel (Corning) coated culture dishes for 1h at 37°C with 5% CO2 using mTeSR1 medium (STEMCELL technologies). Cultures were routinely tested for mycoplasma contamination using MycoAlert Kit (Lonza). Cells were dissociated into small aggregates using ReLeSR (STEMCELL technologies) every 5-7 days and split into freshly coated Matrigel culture dishes.

### Astrocyte generation from NPCs

For the differentiation of astrocytes, a modified protocol was used for NPC generation (Shi et al., 2012). In brief, mycoplasma free iPSCs were seeded in matrigel-coated dishes. To start NPCs generation in 2D, iPSCs were grown until they reached 100% confluency. Optimized neural induction medium was used 1:1 mixture of DMEM/F12 (Gibco) and Neurobasal (Gibco), supplemented with 1:200 N2 supplement (Gibco), 1:100 B27 supplement without vitamin A (Gibco), 50μl 2-mercapthoethanol, 5μg/ml Insulin (Sigma), 1:100 L-Glutamin (Gibco), and 1:100 MEM-NEAA (Gibco), including two SMAD pathway inhibitors 2,5 μM dorsomorphin (Sigma) and, 10μM SB431542 (Selleckchem) to drive differentiation of iPSCs to neural lineage.

The neural induction medium was changed every day, and cells were monitored for the morphological changes during the differentiation time between 8-10 days after plating. When the neuroepithelium sheet was properly formed, then cells were split in aggregates of 300 to 500 cells using Dispase 1mg/ml for 30-45 min at 37°C.

Aggregates were seeded into 6 cm coated PLO and Laminin plate with neural induction medium supplemented with 20ng/ml bFGF (Peprotech) and 10μM Rock inhibitor for 24h. bFGF is used to promote the expansion of NSCs but does not block neural differentiation. The medium was changed the next day with a neural maintenance medium. Low passages NPCs were used to start astrocyte differentiation. NPCs are splitted 24h before, and confluency was maintained to 100% when the medium is changed to astrocyte differentiation medium. Astrocytes differentiation medium contained Neurobasal (Gibco) supplemented with 20ng/ml IGF1 (Peprotech), 10ng/ml Heregulin (Peprotech) 10ng/ml CNTF (Peprotech), 1x Glutamax (Gibco), 1x B27 without vitamin A (Gibco), 100μg/ml Primocin (InvivoGen).

The astrocyte differentiation medium was changed every day until the morphology change was observed. After 15 days, cells were split and analyzed for CD44, an intermediate astrocyte marker. The cells were maintained in the astrocyte differentiation medium until 80 % of the cells expressed GFAP and S100β. Astrocyte growth medium,1:1 mixture of DMEM/F12(Gibco) and Neurobasal (Gibco), EGF 5ng/ml (Peprotech), B27 without vitamin A 1x (Gibco), N2 (Gibco), Primocin 100μg/ml (InvivoGen) was used to culture astrocytes.

### Depletion of IFT88 by siRNA

Synthetic siRNA oligonucleotides were obtained from SMARTPool (Dharmacon). Naïve U3047MG and Nek2-KD expressing U3047MG (1×106) were transfected with 40 nM siRNA targeting IFT88 or with a scrambled sequence (negative control siRNA) for 72h using TransIT-X2 Dynamic (Mirus) transfection kit. The expression of Nek2-KD was induced using doxycycline for another 48h after siRNA IFT88 transfection. The knockdown of IFT88 was assessed by immunofluorescence staining for IFT88. Allstars Negative Control siRNA (Qiagen) was used as a scramble siRNA.

### Immunofluorescence microscopy and imaging of organoids

For live imaging, organoids were grown in 35 mm air diffusing low adherent plate Lumox dish (Sarstedt), which has low autofluorescence and high light transmission properties. Alternatively, we also used μ-Slide angiogenesis slides (Ibidi) for imaging live organoids. Images were acquired using a Leica SP8 laser scanning confocal microscope. Images were captured using 20X air objective. The resulting 8-bit image files were imported into Fiji (ImageJ 1.52i) and maximum intensity projected. Finally, the TIFF files from Fiji were processed using Photoshop (Adobe CC 2018).

Tissue cleared organoids were imaged using Zeiss LSM 880 Airyscan confocal microscope equipped with laser lines 405, 488, 561, and 633 nm, a GaAsP detector, two PMT detectors Plan-Neoflaur 10x/0.3, and Plan-Apochromat 20x/0.8 M27 objectives. The tissue organoids were placed in ECI in 35 mm air diffusing or μ-Slide angiogenesis slides while imaging. 3D image stacks were acquired for representative organoids. The interval between the stacks was kept 2-3 μm apart, depending on the size of the organoids. The captured image files were imported into Fiji. The data were further processed using Image J, Adobe photoshop CC 2018, and Adobe illustrator CC 2018. The z-stack movies were prepared using Fiji. Image-based quantifications were performed manually using Fiji.

### Immunohistochemistry on patient-derived GSCs

GSCs were grown on coverslips fixed in 4% paraformaldehyde (PFA) or ice-cold methanol, for the PFA fixed cells permeabilization was performed with 0.5% Triton X-100 in PBS for 10 min, and then blocked with 0.5% fish gelatin in PBS for 1h at RT. Primary antibody labeling was performed for 1h at RT or overnight at 4°C, followed by three washes in PBS. Primary antibodies used in this study included mouse anti-γ-tubulin (Sigma-Aldrich), rabbit anti-Arl13b (Proteintech), mouse anti-acetylated tubulin (Sigma-Aldrich), rabbit anti-Nde1 (Proteintech), mouse anti-Nestin 4D11 (Novus Biologicals, Cambridge, USA), rabbit anti-Tuj1 (1:100, Sigma-Aldrich), mouse anti-Pax6 (DSHB, Iowa University, Iowa, USA), mouse anti-Nek2 (BD Bioscience), rabbit anti-OFD1 (Gift from Prof. Jeremy Reiter), mouse anti-AuroraA (Cell signaling), mouse anti-HDAC6 (Santa Cruz), mouse anti-CD133 (DSHB), mouse anti-CD15 (DSHB), mouse anti-PDGFR-a (Santa Cruz), rabbit anti-SOX2 (Millipore), mouse anti-FLAG (Sigma), rabbit anti-TUJ1 (Sigma), rabbit anti-GFAP (Cell signaling), rabbit anti-N-cadherin (Abcam), rabbit anti-S100beta (Abcam), mouse anti-GT335 (Adipogen), rabbit anti-IFT88 (Proteintech). The following day, the cells were washed with PBS, incubated with fluorophore-conjugated secondary antibodies 1h at room temperature. Images were acquired using a Leica SP8 scanning confocal microscope and processed using Adobe Photoshop and Illustrator.

STED imaging was performed using TCS SP8 gSTED, Leica. Far-red depletion laser (STED-Laser 775nm) was used for STED imaging. The alignment between channels was monitored using the Gatta STED Nanoruler (Gatta quant, Germany) to monitor STED performance. The PL Apo 100x/1.40 Oil STED Orange (Leica) objective was used, resulting in 100 × overall magnification with ∼50nm lateral and 120nm axial resolution. Signals were detected using gate able hybrid detectors (HyD). Nyquist sampling criteria were maintained during imaging to achieve X-Y resolution of 120nm. Images obtained were deconvoluted using Huygens essential deconvolution software. Line profiles were plotted based on fluorescent intensity values. The images were further processed using ImageJ and Adobe Photoshop.

### GSCs invasion into brain organoid

Brain organoids were generated as previously demonstrated (Gabriel and Gopalakrishnan, 2017; Gabriel et al., 2017). In brief, mycoplasma-free iPSCs were cultured and checked for appropriate stem cell morphology. Once iPSCs reached 80% confluency cells were disassociated into single cells using Accutase (Sigma-Aldrich) for 5 min at 37°C. To start the organoid generation, 35.000 iPSCs were seeded into CellCarrier Spheroid ULA 96-well microplates (PerkinElmer) using neural induction medium (NIM, STEMCELL technologies) containing 10μM ROCK inhibitor Y-27632 (Biozol) for 24h. 100μl of the mix was suspended into cell suspension was given into each well of ULA 96 well U-bottom and incubated at 37°C in the presence of 5% CO2. This process helps in the formation of neurospheres, and the medium is changed once every day for the next 5days. At day-5, neurospheres were embedded a droplet of Matrigel (Corning) using organoid-embedding sheet (STEMCELL technologies). Droplets were solidified at 37°C and were grown four days without agitation in neurosphere medium containing a 1:1 mixture of DMEM/F12 and Neurobasal supplemented with N2 (1:200) (Thermo Fisher Scientific), B27 (1:100) supplement without vitamin A (Thermo Fisher Scientific), 50μl 2-mercapthoethanol, Insulin (Sigma), 1:100 L-Glutamin (Gibco), and 1:100 MEM-NEAA (Gibco). After four days, neurospheres were transferred to a spinner flask containing brain organoid medium which is essentially neurosphere medium supplemented with two SMAD pathway inhibitors 2,5 μM dorsomorphin (Sigma) and 25 μm SB431542 (Selleckchem).

To study GSCs invasion, 10-day old brain organoids were shifted to 35 mm air diffusing low adherent plate Lumox dish (Sarstedt) and 1000 GSCs were provided at the vicinity of the organoids as depicted in the experimental scheme **(Figure 7).** After 48h incubation, organoids with U3047MG Nek2-KD were induced with doxycycline. Organoids supplemented with GSCs before and after cilium induction were subjected to live imaging. The fixed organoids were supplemented with GSCs cells for 10-days, and then doxycycline was supplied for 10-days to induce expression of Nek2-KD. From this, organoids were selected for cryosection and tissue clearing followed by 3D imaging at day-20 after GSCs invasion.

### Whole organoid tissue clearing

Tissue clearing was performed based on the previously described method adapting additional modifications and optimizations (Klingberg et al., 2017). In brief, organoids incubated with GSCs were fixed in 4%PFA (Merck Millipore) for 30 min. Organoids were dehydrated sequentially in graded ethanol (EtOH) series of 30% EtOH (vol/vol), 50% EtOH (vol/vol), 75% EtOH (vol/vol) and 100% EtOH (vol/vol) (each step for up to 5-10 min at 4°C in a gently shaking 2ml tube depending on the size of organoids). Subsequently, tissue clearing was performed with Ethyl cinnamate (ECI; Sigma Aldrich) for approximately 10-20 min at room temperature, depending on the size of organoids. Clarified organoids were then placed into coverslip bottom μ-slides (Ibidi) and stored at 4°C until imaging.

### Intracranial xenografts of human GSCs in mouse

Experiments involving animals were approved by the Ethical Committee of the Istituto Superiore di Sanità, Rome (Pr. No. 4701/17). Immunosuppressed NOD SCID mice (male, 20-23 g; Charles River, Milan, Italy) were anesthetized with intraperitoneal injection of diazepam (2 mg/100 g) followed by intramuscular injection of ketamine (4 mg/100 g). Animal skulls were immobilized in a stereotactic head frame and a burr hole was made 2 mm right of the midline and 1 mm anterior to the bregma. The tip of a 10 μl-Hamilton microsyringe was placed at a depth of 3 mm from the dura and 5 μl of PBS containing 2×10^4^ of either mCherry expressing U3047MG cells were slowly injected (D’Alessandris et al., 2017) (Ricci-Vitiani et al., 2010). The U3047MG cells were engineered to express catalytically inactive Nek2-KD upon doxycycline induction. Solutions containing 1 mg/ml doxycycline hyclate (Sigma-Aldrich, St Louis, MO, USA) and 1% (w/v) sucrose were prepared into tap water and protected from light during experiments. After grafting, the animals were kept under pathogen-free conditions in positive-pressure cabinets (Tecniplast Gazzada, Varese, Italy) and observed daily for neurological signs and body weight. After survivals ranging from 4 to 24 weeks, the mice were deeply anesthetized and transcardially perfused with 0.1 M PBS (pH 7.4) then treated with 4% paraformaldehyde in 0.1 M PBS. The brain was removed and stored in 30% sucrose in PBS for 3 days.

### Fluorescence microscopy and immunofluorescence of brain tumor xenografts

Brains were frozen sectioned on the coronal plane (20 μm thick). For fluorescence microscopy, sections 120 μm apart were collected in distilled water and mounted on slides with Vectashield mounting medium (Bio-Optica, Milan, Italy). Images were acquired with a laser scanning confocal microscope (LSM 500 META, Zeiss, Milan, Italy). The cranio-caudal extension of the brain area invaded by fluorescent tumor cells was assessed on serial coronal sections.

For immunofluorescence, slices were rinsed three times at room temperature (10 min each) in PBS, and then blocked in PBS with 10% BSA, 0.3% Triton X-1000 for 2 hours. Sections were then incubated overnight at 4°C in PBS with 0.3% Triton X-1000, 0.1% normal donkey serum (NDS) with primary antibodies. For Arl13B detection, slices were incubated with the following primary antibodies: rabbit anti-Arl13B (1:200, Proteintech) or mouse anti-ARL13B (1:500, NeuroMab, Davis, CA, USA). Slices were then rinsed three times in PBS (10 min each) at room temperature and incubated in PB containing 0.3% Triton X-100 with secondary antibodies for 2 hours at RT. Secondary antibodies used were as follows: Alexa Fluor 647 or 555 donkey anti-mouse, and Alexa Fluor 555 or 647 donkey anti-rabbit secondary antibodies (1:500; Thermo Fisher Scientific, Waltham, MA).

### Immunohistochemistry on human specimens

All patients provided written informed consent to the study according to research proposals approved by the Institutional Ethics Committee of Fondazione Policlinico Gemelli, UCSC (Prot. 4720/17). Tissue samples were fixed in 4.5% formalin for 48 hours at 4°C, post-fixed in 30% sucrose, and sectioned (40 μm) by a cryostat. The sections were permeabilized overnight at 4°C in PBS with 0.3% Triton X-and 0.1% NDS with primary antibodies. Primary antibodies, rat anti-Collapsin Response-Mediated Protein 5 (CRMP5, 1:50, Millipore), mouse anti-γ-tubulin (1:50, Santa Cruz), rabbit anti-ARL13B (1:200, Proteintech). Following washed slices were incubated with fluorophore-conjugated secondary for 2h at room temperature. Images were acquired with a laser confocal microscope (Leica SP5).

### Electron microscopy

GSCs were grown on coverslips and fixed with 2% glutaraldehyde (Electron Microscopy Sciences), and processed in electron microscopy as previously described (Gopalakrishnan et al., 2012; Gopalakrishnan et al., 2011). The embedded cell pellets were ultra-thin-sectioned (80 nm), counterstained, and visualized for ultra-structural details of centrosomes and cilia using a Zeiss 10A electron microscope. Images were processed using Adobe Photoshop.

### Western blotting

Total protein from cultured cells was homogenized in RIPA lysis buffer with a mixture of protease and phosphatase inhibitors (Santa Cruz). Protein concentration was determined using the Bradford assay (Sigma-Aldrich), and samples were run on a 10% SDS-page gel. Primary antibodies used were mouse anti-CPAP (Hybridoma C44), mouse anti-Nek2 (BD Bioscience), rabbit anti-ODF2 (Proteintech), rabbit anti-GAPDH (Proteintech), rabbit anti-OFD1 (Gift from Prof. Jeremy Reiter), rabbit anti-Nde1 (Proteintech), mouse anti-AuroraA (Cell signaling), rabbit anti-Plk1 (Cell signaling), mouse anti-HDAC6 (Santa Cruz), mouse anti-GFP (Proteintech), rabbit anti-myc (Abcam), mouse anti-CD133 (DSHB), mouse anti-PDGFR-a (Santa Cruz), rabbit anti-AKT (Cell signaling), rabbit anti-phopho-AKT (Cell Signaling), rabbit anti-SOX2 (Millipore), mouse anti-FLAG (Sigma), rabbit anti-TUJ1 (Sigma), rabbit anti-GFAP (Cell signaling). Membranes were probed with horseradish peroxidase-linked donkey anti-rat IgG (1:5000), goat anti-mouse IgG (1:5000) and detection of protein was conducted using ECL Western blotting reagents (Thermo Fisher). All Western blots are representative images from at least three biological replicates.

### FLAG-immunoprecipitation (IP) for CPAP and Nek2 complexes

Flag coted beads (Sigma Aldrich) were incubated with high-speed lysate (HSL) for 3h at 4°C. The lysates were prepared using HEK293T cells stable expressing pSIN CPAP-WT 3xflag and pLenti Nek2-WT 3xflag lysed with 1x BRB80 buffer (extract buffer) as described previously (Gopalakrishnan et al., 2011). After incubation, the beads were washed 3x times with extract buffer and washed 2x with high-salt buffer containing 500 mM salt, and final wash with buffer containing 100 mM NaCl, the samples were eluted by boiling them in 2x Laemmli buffer at 98°C. The beads were analyzed by western blotting for the CPAP and Nek2-complexes.

### RNA sequencing and analysis

U3047MG (before ciliation) and U3047-Nek2-KD (after ciliation) cells were collected and washed once with cold PBS before adding to 300 μl Tri-Reagent (Sigma Aldrich, USA) by free-thawing, and total RNA was isolated and DNase-treated using the DirectZol RNA kit (Zymo Research). Approximately 2 μg of total RNA was used to subselect poly(A)+ transcripts and generate strand-specific cDNA libraries (TrueSeq kit; Illumina).

Poly(A)-enriched RNA was prepared and sequenced on a HiSeq4000 platform (Illumina; strand-specific) to >35×106 reads per sample. Reads were quality assessed and mapped to hg19 using STAR (Dobin et al., 2013), followed by quantification of unique exon counts using featureCounts (Liao et al., 2014). Counts were further normalized via the RUVs function of RUVseq (Risso et al., 2014) to estimate factors of unwanted variation using those genes in the replicates for which the covariates of interest remain constant and correct for unwanted variation, before differential gene expression was estimated using DESeq2 (Love et al., 2014). Genes with an FDR <0.01 and an absolute (log2) fold-change of >0.6 were deemed differentially-expressed and listed in Table S1

### Statistical analysis

Data were analyses statistically using GraphPad Prism 7 for Mac OS X (Version 7.0e, September 5, 2018). Data are presented as mean ± S.E.M. The following tests were used to obtain P values: two-way ANOVA, followed by Sidak’s multiple comparisons test or Tukey’s multiple comparisons test or unpaired t test or Dunnett’s multiple comparisons test. n.s, indicates no significance, *P<0.01, **P<0.001, ***P<0.0001.

